# Functional 3’-UTR Variants Identify Regulatory Mechanisms Impacting Alcohol Use Disorder and Related Traits

**DOI:** 10.1101/2024.01.31.578270

**Authors:** Andy B. Chen, Xuhong Yu, Kriti S. Thapa, Hongyu Gao, Jill L Reiter, Xiaoling Xuei, Andy P. Tsai, Gary E. Landreth, Dongbing Lai, Yue Wang, Tatiana M. Foroud, Jay A. Tischfield, Howard J. Edenberg, Yunlong Liu

## Abstract

Although genome-wide association studies (GWAS) have identified loci associated with alcohol consumption and alcohol use disorder (AUD), they do not identify which variants are functional. To approach this, we evaluated the impact of variants in 3’ untranslated regions (3’-UTRs) of genes in loci associated with substance use and neurological disorders using a massively parallel reporter assay (MPRA) in neuroblastoma and microglia cells. Functionally impactful variants explained a higher proportion of heritability of alcohol traits than non-functional variants. We identified genes whose 3’-UTR activities are associated with AUD and alcohol consumption by combining variant effects from MPRA with GWAS results. We examined their effects by evaluating gene expression after CRISPR inhibition of neuronal cells and stratifying brain tissue samples by MPRA-derived 3’-UTR activity. A pathway analysis of differentially expressed genes identified inflammation response pathways. These analyses suggest that variation in response to inflammation contributes to the propensity to increase alcohol consumption.

## Introduction

Genome-wide association studies (GWAS) have identified loci associated with alcohol use disorder^1–4^ (AUD), alcohol consumption^1,2,5–9^, and related traits^10–18^. Though these strategies have expanded our understanding of the genetics of alcohol related traits, they are still limited in how much they can unveil about the underlying mechanisms. Variants in protein coding regions of the genome may cause a change in protein function, but most significant variants occur in noncoding regions and may be genetic markers rather than have a functional effect. To address this, massively parallel reporter assays (MPRAs) have been developed to identify variants in the broad loci that alter gene expression^19,20^. In this study, we used PASSPORT-seq^21,22^ to evaluate variants in the 3’ untranslated regions (3’-UTR) of genes within those loci.

Though MPRAs can evaluate the effect of candidate variants, they cannot elucidate their roles in disease. Several methods use a Mendelian Randomization (MR)-like framework to identify genes that contribute to the onset of a phenotype, including PrediXcan^23^, TWAS^24^, and SMR^25^. These methods impute gene transcription from genotype using models trained on tissue-level transcriptome datasets, and test the association of these imputed gene expression levels with the phenotype. By using imputed expression, these approaches seek to isolate the gene’s effect on the phenotype while removing the phenotype’s effect on gene expression. These imputations, however, are based on association of variants with gene expression rather than the actual functional effect of each variant. To address this, we propose MPRA-mediated Gene Expression Association Analysis (MGExA), an approach that combines variant function derived from MPRA with GWAS summary statistics, to identify genes whose imputed expression based on genotype and MPRA results is associated with the phenotype. Our results suggest that this strategy not only provides increased statistical power for discovering genes that contribute to AUD and related traits, but also enables identifying which cell types those genes function in.

## Results & Discussion

### MPRA identifies functional variants impacting gene expression

To identify genes within GWAS loci likely to functionally impact AUD and related traits, we started with a set of SNPs that were weakly associated (p < 10^−5^) with AUD, alcohol-related traits, and neurological disorders (Figure S1, Table S1 and S2; ref. ^26,27^). We then selected 3’-UTR SNPs within regions of LD (r^2^>0.8) around the initial set if they had a minor allele frequency greater than 5% in at least one of the five super-populations from phase 3 of the 1000 Genomes Project^28^. This resulted in a total of 13,515 candidate SNPs (see Methods and Figure S1), from which we created a pool of 24,780 oligos (some SNPs shared a reference sequence), each extending 25 base pairs upstream and downstream from both the reference and alternative alleles of each candidate SNP. This pool was cloned into the pIS-0 vector (Methods; Figure S2A) and transfected 6 independent times into two human cell lines, SH-SY5Y neuroblastoma cells^29^ and SV-40-immortalized microglial cells (Cat.No: T0251, Applied Biological Materials Inc, Richmond, QC, Canada). After 42 h, the cells were harvested, the DNA and RNA were isolated, cDNA synthesized, and sequencing libraries prepared that included barcodes and unique molecular indices (Figure S2B). The number of unique cDNA and DNA reads of reference and alternative alleles for each oligo was determined by sequencing (Figure S3).

For each oligo, we compared the normalized number of DNA and RNA (cDNA) reads using a generalized linear model (see Methods) to determine if the oligo altered gene expression. If the inserted 3’-UTR sequences have no effect on transcription, the RNA abundance should be proportional to the DNA abundance. Of the 24,780 oligo sequences, 2,908 altered gene expression (FDR < 0.05) in SH-SY5Y; of those, 81 increased expression by more than 2-fold, while 876 decreased expression by more than 2-fold (Figure 1A, Table S3). In microglia cells, 5,460 sequences altered gene expression (FDR < 0.05), with 71 increasing expression by more than 2-fold and 1,220 decreasing expression by more than 2-fold. This is consistent with the expectation that 3’-UTR sequences are enriched with *cis*-acting elements that regulate RNA degradation.

**Figure 1.**
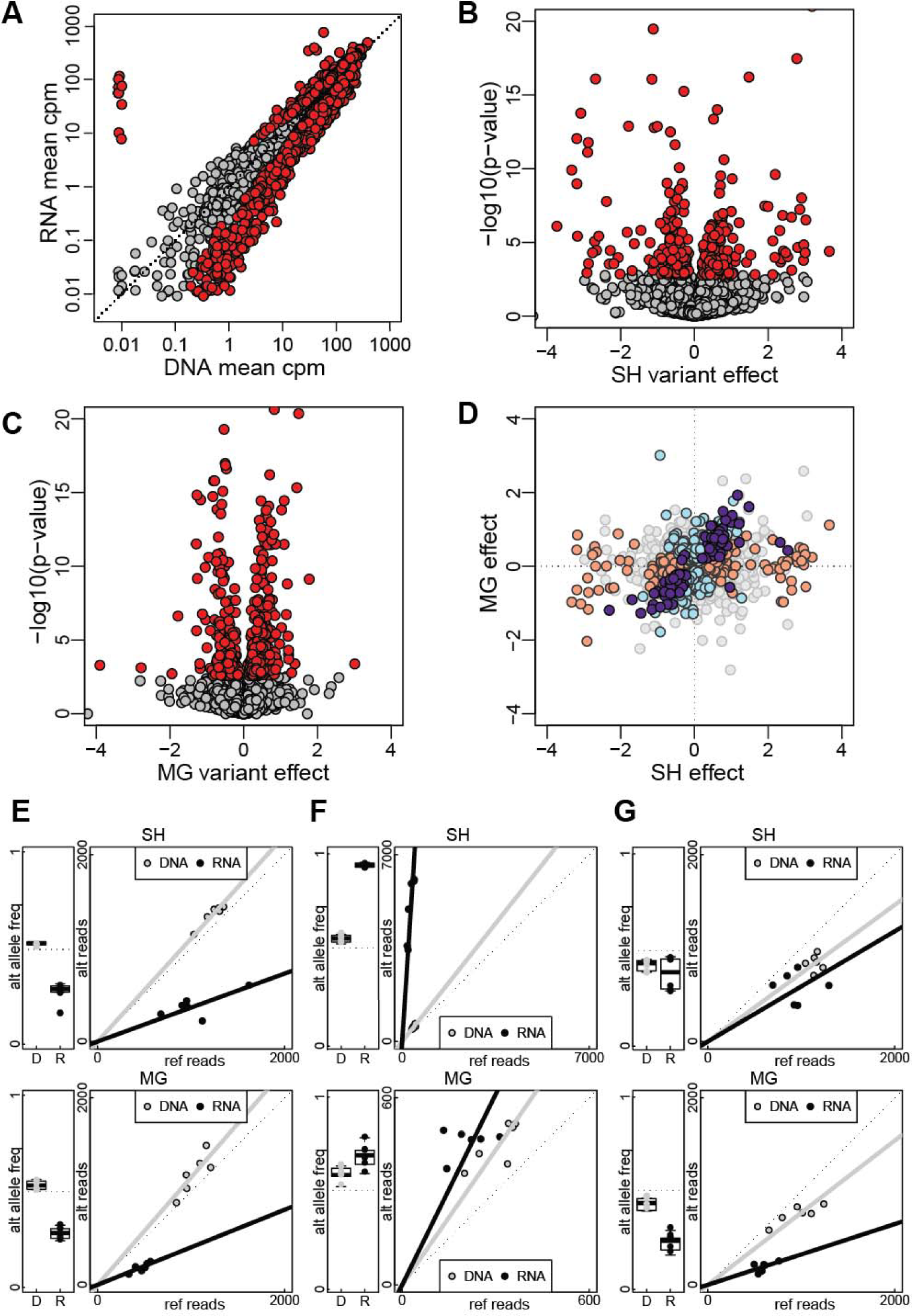
Variant effects on gene expression. (A) Scatterplot comparing the mean counts per million reads (CPM) of RNA (cDNA) and DNA for each oligonucleotide following transfection in SH-SY5Y cells. Red dots denote oligos where the RNA and DNA mean CPM were significantly different (FDR < 0.05). CPM was calculated by dividing the oligonucleotide count by the total number of reads in that sample multiplied by 10^6^. (B) Effects of SNPs on gene expression in SH-SY5Y cells. Red dots denote SNPs for which the variant significantly (FDR < 0.05) affected gene expression. (C) Effects of SNPs on gene expression in microglia cells. Red dots denote SNPs for which the variant significantly (FDR < 0.05) affected gene expression. (D) Comparison of the effect of SNPs in SH-SY5Y (SH) and microglia (MG) cell lines. Light orange denotes SNPs significant (FDR < 0.05) in SH-SY5Y cells, light blue denotes those significant in microglia cells, and dark purple denotes SNPs significant in both cell lines. Gray dots represent SNPs that were not significant in either cell line. (E-G) Examples of assay results in SH-SY5Y and microglia cells for 3 different SNPs: rs2298753 on ADH1C (E), rs1139697 on PDLIM5 (F), and rs4736367 on JRK (G). Boxplot of alternative allele frequency in DNA and RNA and scatterplot of reference and alternative counts for RNA and DNA.

We then used a generalized linear mixed effect model to determine whether the alternative alleles differentially affected gene expression. Of the 13,515 SNPs tested, 972 (7.2%) showed significant differences between alleles (FDR <0.05): 400 in SH-SY5Y cells and 657 in microglia cells (Figure 1B & C, Table S3). Among these, only 84 SNPs showed significant allelic imbalance in both cell lines, 81 of which affected expression in the same direction (Figure 1D).

### Functionally impactful variants explained more heritability than expected

We tested whether the 3’-UTR SNPs that differentially affected gene expression explained a disproportionate fraction of heritability for the number of alcoholic drinks per week (DPW; ref ^6^) or alcohol use disorder (AUD; ref. 2). For this, we analyzed the subset of SNPs for which we had both MPRA data and GWAS data. For DPW^6^, 223 SNPs among the 6,080 in SH-SY5Y and 379 among 6,119 in microglia significantly affected gene expression in our MPRA. These SNPs explained 1.4% and 2.3% of the overall heritability, respectively. This was 2.1-fold (SH-SY5Y; *p* = 0.034) and 2.3-fold (microglia; *p* = 0.003) as much as the median of 1,000 randomly selected sets of equal size chosen from among all the tested SNPs (Table S4 and Figure 2). Similarly, for AUD the 158 (among 4,348 tested; SH-SY5Y) and 285 (among 1,306; microglia) functional SNPs explained 1.5% (SH-SY5Y) and 1.3% (microglial) of the heritability, which is 8.4 times (p=0.001) and 5.0 times (p=0.004) the median heritability of an equal-sized randomly selected set of the tested candidate MPRA SNPs (Table S4).

**Figure 2.**
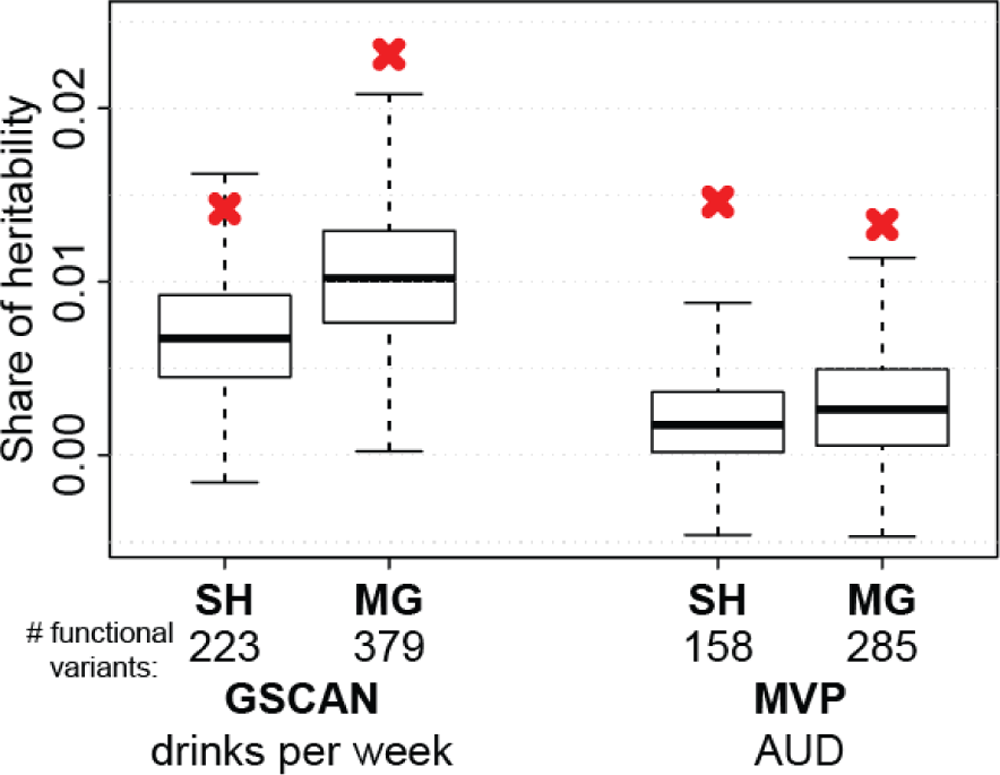
Heritability enrichment of the sets of SNPs with significant effects found by MPRA in SH-SY5Y (SH) and microglia (MG) cells. Heritability enrichment was evaluated in GWAS of alcoholic drinks per week, from GSCAN, and alcohol use disorder (AUD), from the MVP. Red X marks show the heritability of the SNPs shown by MPRA to be functional in each cell line, with the number of functional variants shown below. Results of the permutation tests of 1000 random sets of equal size from among all candidate SNPs are shown as boxplots.

### Combining GWAS and MPRA identifies genes whose 3’-UTR activity is associated with alcohol consumption and AUD

To examine how the genetic variants in the non-coding regulatory regions of a gene might contribute to a trait of interest, we designed a computational framework, which we call MPRA-mediated Gene Expression Association Analysis (MGExA). MGExA multiplies the functional effect of each SNP (determined by the MPRA) by its effect size from a GWAS of a trait and sums the results for each gene to calculate a Z-score (Wald statistic *U_m_*) for the association between the trait and the imputed gene expression levels (Methods, equation 3, and Figure S4A)^2,6^. We used MGExA to identify a set of genes whose expression levels, as calculated based upon the 3’-UTR genotypes, are associated with DPW^6^. We replicated our finding using the summary statistics from the AUDIT-C GWAS in the Million Veteran Program (MVP) study^2^. Although DPW and AUDIT-C characterize different aspects of drinking behavior, they are both related to alcohol consumption.

Of the genes that contained at least one MPRA-evaluated 3’-UTR variant with *p* < 0.05 in the GSCAN study (Table 1, Table S5), 3’-UTR activities of 38 genes in SH-SY5Y (out of 596) and 50 genes in microglia (out of 690) were significantly associated with DPW (FDR < 0.2); 12 were common to both cell lines. Among these, 17/31 genes in SH-SY5Y cells (7 could not be tested because of differences in the variants available between datasets) and 14/45 in microglia (5 not tested) were also significantly associated with AUDIT-C (FDR<0.2); four of these were significant in both cell types (Figure 3A & B). The direction of the effect in 16 out of the 17 genes identified based on MPRA in the SH-SY5Y cells and in all 14 genes from microglia was consistent across both studies (Table 1; Figure 3A & B).

**Figure 3.**
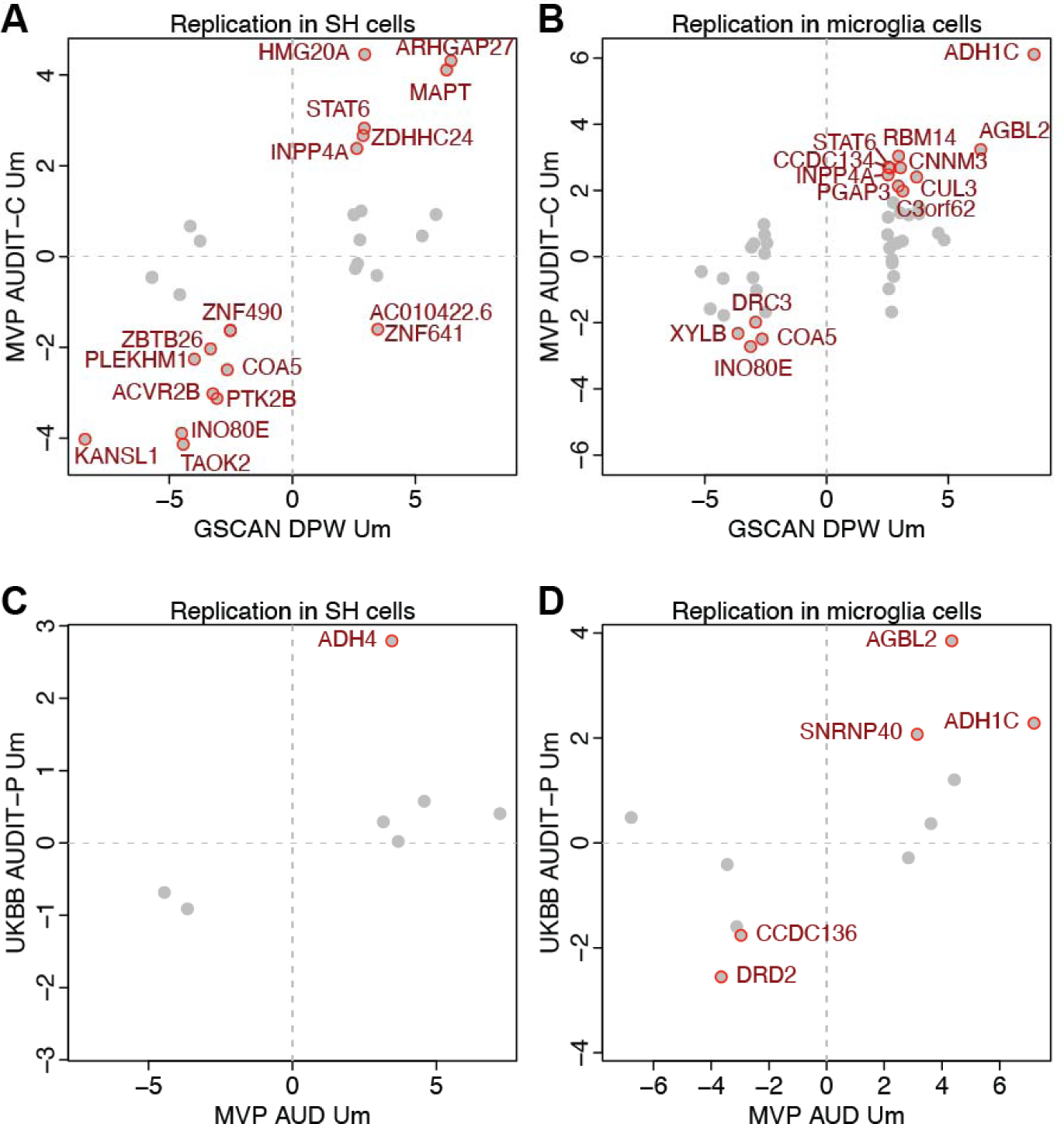
Gene identification using MPRA-mediated Gene Expression Association (MGExA). Scatterplots of the results of applying MGExA to identify genes whose aggregated 3’-UTR variants are associated with alcohol consumption and alcohol use disorder. Plots compare the MGExA results between drinks per week^6^ and AUDIT-C^2^ in (A) SH-SY5Y and (B) microglia cells and between AUD^2^ and AUDIT-P^1^ in (C) SH-SY5Y and (D) microglia cells. The *U_m_* values for genes significant (FDR < 0.2) in discovery studies (DPW and AUD) are plotted in gray; and those replicated (AUDIT-C and AUDIT-P) (FDR < 0.2) were labeled and outlined in red.

**Table 1.** Genes identified by MGExA as having 3’-UTRs associated with alcohol consumption or AUD. Genes were first identified by MGExA using GWAS summary statistics of drinks per week^6^ phenotype or AUD, and genes with FDR < 0.2 were then evaluated by MGExA using AUDIT-C or AUDIT-P. Table contains genes with FDR < 0.2 in both the discovery (drinks per week and AUD) and replication (AUDIT-C and AUDIT-P) studies. Genes identified in both SH-SY5Y and microglia cell lines are starred (*). Um: the statistic derived from MGExA; p: the p-value associated with Um; FDR: the false discovery rate of p; SNPs: the number of SNPs used to evaluate each gene.

|  |  | discovery study |  |  |  | replication study |  |  |  |
| --- | --- | --- | --- | --- | --- | --- | --- | --- | --- |
| gene | gene name | Um | p | FDR | SNPs | Um | p | FDR | SNPs |
| Alcohol Consumption: SH-SY5Y |  |  |  |  |  |  |  |  |  |
| ARHGAP27 | Rho GTPase activating protein 27 | 3.8 | 1.58E-04 | 6.73E-03 | 7 | 3.2 | 1.46E-03 | 8.75E-03 | 5 |
| CHADL* | chondroadherin like | 2.8 | 4.61E-03 | 9.76E-02 | 3 | 2.7 | 7.15E-03 | 2.73E-02 | 3 |
| COA5* | cytochrome c oxidase assembly factor 5 | -2.6 | 8.61E-03 | 1.47E-01 | 2 | -2.5 | 1.34E-02 | 3.76E-02 | 2 |
| DBIL5P2 | diazepam binding inhibitor-like 5 pseudogene 2 | 3.7 | 2.54E-04 | 9.47E-03 | 3 | 2.5 | 1.12E-02 | 3.35E-02 | 2 |
| HMG20A | high mobility group 20A | 2.9 | 3.31E-03 | 7.88E-02 | 1 | 4.5 | 8.48E-06 | 3.56E-04 | 1 |
| INO80E* | INO80 complex subunit E | -4.2 | 2.59E-05 | 1.71E-03 | 3 | -3.6 | 2.64E-04 | 1.85E-03 | 2 |
| INPP4A* | inositol polyphosphate-4-phosphatase type I A | 3.1 | 1.66E-03 | 4.70E-02 | 9 | 2.5 | 1.09E-02 | 3.35E-02 | 8 |
| KANSL1 | KAT8 regulatory NSL complex subunit 1 | -5.3 | 9.87E-08 | 1.47E-05 | 8 | -4.0 | 6.45E-05 | 6.77E-04 | 11 |
| LINC02210-CRHR1 | LINC02210-CRHR1 readthrough | 8.6 | 0.00E+00 | 0.00E+00 | 11 | 4.1 | 3.90E-05 | 5.46E-04 | 11 |
| MAPT | microtubule associated protein tau | 6.7 | 1.75E-11 | 5.22E-09 | 14 | 3.9 | 1.03E-04 | 8.61E-04 | 14 |
| PTK2B* | protein tyrosine kinase 2 beta | -3.2 | 1.59E-03 | 4.70E-02 | 2 | -2.9 | 3.89E-03 | 2.04E-02 | 2 |
| SREBF2 | sterol regulatory element binding transcription factor 2 | 2.9 | 4.09E-03 | 9.02E-02 | 4 | 2.8 | 5.18E-03 | 2.18E-02 | 2 |
| STAT6* | signal transducer and activator of transcription 6 | 3.0 | 2.86E-03 | 7.40E-02 | 6 | 2.8 | 4.53E-03 | 2.11E-02 | 4 |
| TAOK2 | TAO kinase 2 | -4.6 | 4.56E-06 | 3.88E-04 | 2 | -4.2 | 2.61E-05 | 5.46E-04 | 1 |
| TFAP2B | transcription factor AP-2 beta | -2.8 | 5.03E-03 | 1.00E-01 | 12 | -2.2 | 2.83E-02 | 7.43E-02 | 9 |
| WDR82 | WD repeat domain 82 | 2.9 | 3.23E-03 | 7.88E-02 | 5 | -2.7 | 7.85E-03 | 2.75E-02 | 4 |
| ZBTB26 | zinc finger and BTB domain containing 26 | -3.3 | 8.63E-04 | 2.71E-02 | 1 | -2.0 | 4.20E-02 | 1.04E-01 | 1 |
| Alcohol Consumption: microglia |  |  |  |  |  |  |  |  |  |
| ACVR2B | activin A receptor type 2B | 3.7 | 1.86E-04 | 1.29E-02 | 16 | 2.2 | 2.80E-02 | 1.10E-01 | 15 |
| ADH1C | alcohol dehydrogenase 1C (class I), gamma polypeptide | 8.5 | 0.00E+00 | 0.00E+00 | 1 | 6.1 | 9.83E-10 | 4.62E-08 | 1 |
| AGBL2 | AGBL carboxypeptidase 2 | 6.1 | 1.03E-09 | 2.36E-07 | 3 | 2.9 | 3.57E-03 | 3.22E-02 | 2 |
| C3orf62 | chromosome 3 open reading frame 62 | 3.1 | 2.18E-03 | 8.37E-02 | 2 | 1.9 | 5.61E-02 | 1.65E-01 | 2 |
| CHADL* | chondroadherin like | 2.7 | 7.37E-03 | 1.43E-01 | 3 | 2.1 | 4.00E-02 | 1.34E-01 | 3 |
| COA5* | cytochrome c oxidase assembly factor 5 | -2.7 | 6.80E-03 | 1.40E-01 | 2 | -2.5 | 1.12E-02 | 4.78E-02 | 2 |
| CUL3 | cullin 3 | 3.4 | 7.62E-04 | 3.75E-02 | 11 | 2.8 | 4.40E-03 | 3.22E-02 | 10 |
| DRC3 | dynein regulatory complex subunit 3 | -2.9 | 3.38E-03 | 8.41E-02 | 3 | -2.0 | 4.66E-02 | 1.46E-01 | 3 |
| DRD2 | dopamine receptor D2 | -6.4 | 1.52E-10 | 5.23E-08 | 3 | -3.8 | 1.41E-04 | 3.31E-03 | 4 |
| INO80E* | INO80 complex subunit E | -3.8 | 1.57E-04 | 1.26E-02 | 3 | -3.3 | 1.03E-03 | 1.62E-02 | 2 |
| INPP4A* | inositol polyphosphate-4-phosphatase type I A | 3.2 | 1.27E-03 | 5.48E-02 | 10 | 2.8 | 5.89E-03 | 3.22E-02 | 8 |
| MTCH2 | mitochondrial carrier 2 | 5.2 | 2.04E-07 | 3.52E-05 | 12 | 2.7 | 6.17E-03 | 3.22E-02 | 5 |
| PGAP3 | post-GPI attachment to proteins phospholipase 3 | 3.0 | 2.82E-03 | 8.41E-02 | 4 | 2.2 | 3.15E-02 | 1.14E-01 | 4 |
| PTK2B* | protein tyrosine kinase 2 beta | -3.0 | 2.59E-03 | 8.41E-02 | 2 | -1.8 | 7.48E-02 | 1.95E-01 | 2 |
| RBM14-RBM4 | RBM14-RBM4 readthrough | 2.9 | 3.27E-03 | 8.41E-02 | 1 | 3.1 | 1.76E-03 | 2.07E-02 | 2 |
| STAT6* | signal transducer and activator of transcription 6 | 2.6 | 1.05E-02 | 1.69E-01 | 5 | 2.8 | 5.77E-03 | 3.22E-02 | 3 |
| TRIM27 | tripartite motif containing 27 | 2.9 | 3.41E-03 | 8.41E-02 | 2 | 1.8 | 6.67E-02 | 1.84E-01 | 2 |
| XYLB | xylulokinase | -3.4 | 5.84E-04 | 3.10E-02 | 11 | -2.6 | 9.54E-03 | 4.48E-02 | 11 |
| Alcohol use disorder: SH-SY5Y |  |  |  |  |  |  |  |  |  |
| ADH4* | alcohol dehydrogenase 4 (class II), pi polypeptide | 3.5 | 4.06E-04 | 2.98E-02 | 5 | 2.9 | 4.18E-03 | 2.93E-02 | 5 |
| Alcohol use disorder: microglia |  |  |  |  |  |  |  |  |  |
| ADH1C | alcohol dehydrogenase 1C (class I), gamma polypeptide | 7.2 | 6.32E-13 | 3.28E-10 | 1 | 2.3 | 2.25E-02 | 7.88E-02 | 1 |
| ADH4* | alcohol dehydrogenase 4 (class II), pi polypeptide | 3.0 | 2.31E-03 | 9.97E-02 | 4 | 2.4 | 1.52E-02 | 7.11E-02 | 4 |
| AGBL2 | AGBL carboxypeptidase 2 | 4.2 | 2.79E-05 | 3.62E-03 | 2 | 3.4 | 5.62E-04 | 3.93E-03 | 2 |
| CCDC136 | coiled-coil domain containing 136 | -3.0 | 2.94E-03 | 1.17E-01 | 2 | -1.7 | 8.13E-02 | 1.90E-01 | 2 |
| DRD2 | dopamine receptor D2 | -4.2 | 2.64E-05 | 3.62E-03 | 4 | -3.5 | 4.22E-04 | 3.93E-03 | 3 |
| SNRNP40 | small nuclear ribonucleoprotein U5 subunit 40 | 3.1 | 1.68E-03 | 7.90E-02 | 1 | 2.1 | 3.84E-02 | 1.08E-01 | 1 |

We also used MGExA to analyze traits related to AUD: the MVP GWAS for AUD^2^ was the discovery study and AUDIT-P (questions 4-10; related to AUD^1,16^) in the UK BioBank^1^ was the replication study. Of the 441 genes with at least one MPRA-evaluated SNP at *p*<0.05 in SH-SY5Y cells, 3’-UTR activities of 7 genes were significantly associated with AUD (FDR < 0.2), and 1 was replicated using AUDIT-P (Table 1; Figure 3C & D). Of the eligible genes in microglia, 11 out of 519 were significant (FDR < 0.2) with AUD, 5 of which were also significant for AUDIT-P. All significant genes were directionally consistent across the two studies (Figure 3).

Combining GWAS and MPRA data using MGExA led to identification of 31 genes whose 3’-UTR activities contribute to either alcohol consumption or alcohol use disorder. Interestingly, only 5 among these (16%) contain 3’-UTR SNPs that were genome-wide significant (p < 5×10^−8^) in either GWAS^2,6^. The increased statistical power is in part the result of combining the effects of multiple functional variants affecting the same gene with GWAS summary statistics. For example, the weighted contributions of each SNP to the trait and the effect size in the MPRA and GWAS analysis for Diazepam Binding Inhibitor-Like 5 Pseudogene 2 (*DBIL5P2*), TAO Kinase 2 (*TAOK2*), INO80 Complex Subunit E (*INO80E*), and Dopamine Receptor D2 (*DRD2*) are depicted in Figure S4B&C. Each gene’s 3’-UTR contains multiple SNPs evaluated in both the MPRA and GWAS for drinks per week. The combined effects suggest that their gene expression levels positively (for *DBIL5P2*) or negatively (for *TAOK2*, *INO80E*, and *DRD2*) associate with the phenotype. None of these SNPs nor any SNPs in LD with them (r^2^ > 0.8) had GWAS signals that reach genome-wide significance.

Of the genes identified by MGExA as associated with alcohol consumption or alcohol use disorder, 29 are expressed in microglia or neurons in the single cell transcriptome collated in the Human Brain Cell Atlas^30^. Several, including Alcohol Dehydrogenase 4^31^ (*ADH4*), AGBL Carboxypeptidase 2^32^ (*AGBL2*), and Microtubule Associated Protein Tau^33^ (*MAPT*), have previously been shown to be associated with alcohol-related phenotypes. Other MGExA-identified genes, such as Signal Transducer And Activator Of Transcription 6^34^ (*STAT6*) and *TAOK2*^35^ have been found to be involved in the response to ethanol in animal models. MGExA also identified genes associated with addiction to other substances, including nicotine (KAT8 Regulatory NSL Complex Subunit 1^36^; *KANSL1*) and opioids (*DRD2*^37^). MGExA genes were also associated with other disorders of the brain such as Alzheimer’s disease (Protein Tyrosine Kinase 2 Beta^38^; *PTK2B*), and schizophrenia (*SREBF2*^39^).

### CRISPR inhibition of target genes identified potential mechanisms

To identify the downstream pathways of the genes identified by MGExA, we used a KRAB-dCas9-based CRISPRi assay followed by single-cell RNA-seq (Perturb-seq^40^) to measure global gene expression changes after knocking down the target genes in SH-SY5Y cells. We designed 5 single-guide RNAs (sgRNAs) targeting each of the 31 candidate genes whose 3’-UTR variant-determined gene expression levels are associated with alcohol consumption and AUD traits in the discovery studies and replicated. In addition to the 155 candidate sgRNAs, 18 positive control sgRNAs and 18 negative template control sgRNAs were included (Table S6). Lentivirus containing the 191 sgRNAs were transduced into a modified SH-SY5Y cell line that stably expresses KRAB-dCas9 (see Methods). After 8 days of transfection, 10XGenomics single cell RNA-seq assays were conducted; feature barcoding was used to identify the sgRNA transduced into each cell. We found that 8,842 cells were transduced with one target gene sgRNA, and 1,331 cells were transduced with a negative template control. Using Seurat Mixscape^41^, seven genes were found to have robust perturbation responses: *CUL3, INO80E, KANSL1, PLEKHM1, RBM14, SNRNP40*, and *TAOK2* (Figure 4A&B). For each of these genes, we evaluated differential gene expression between perturbed cells compared with non-targeting control (NTC) cells using the Wilcoxon Rank-Sum test. Four target genes had more than 10 significant (adjusted p < 0.05) differentially expressed genes: *CUL3* (17), *KANSL1* (199), *RBM14* (505), and *SNRNP40* (49) (Table S7).

**Figure 4.**
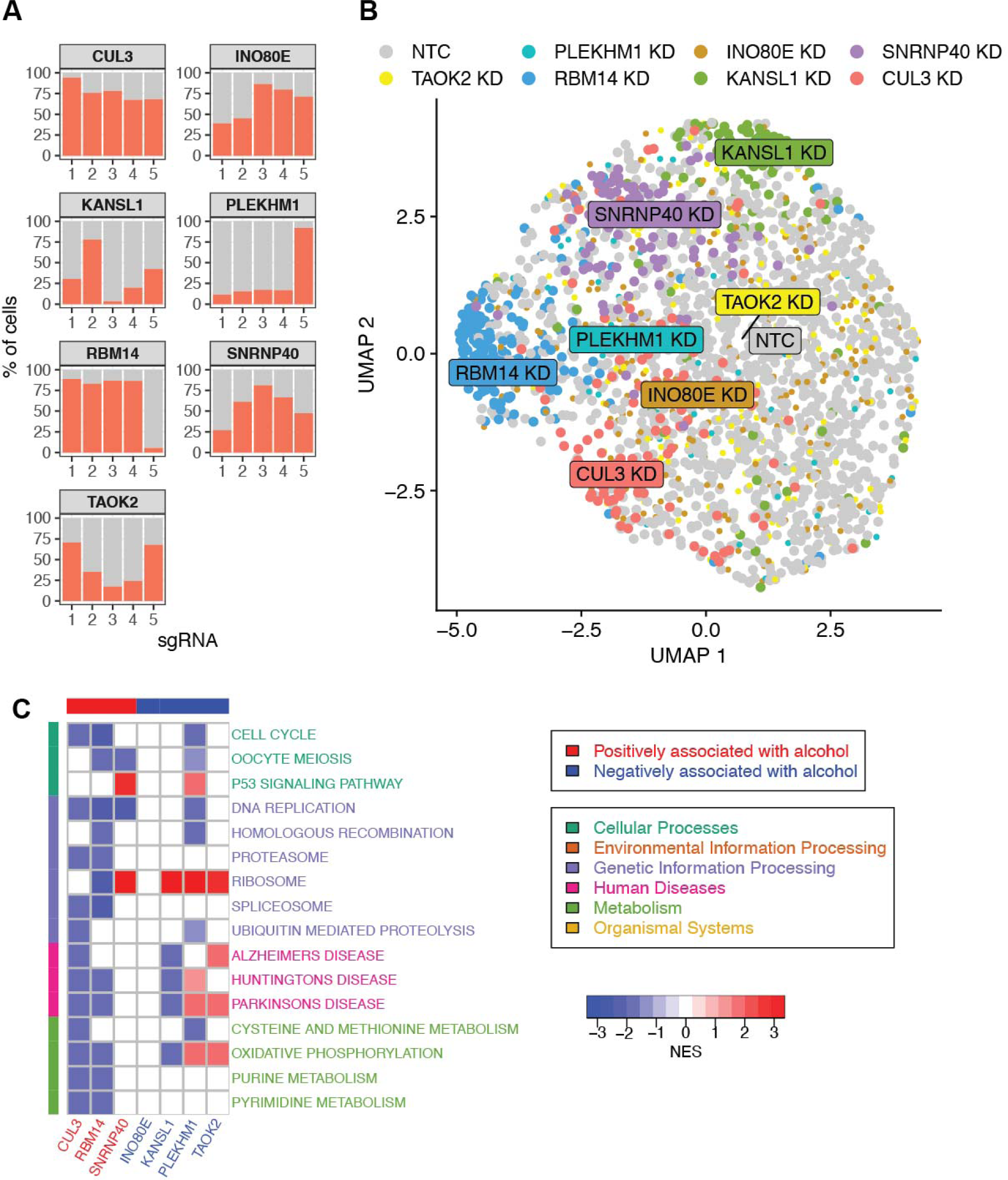
CRISPRi-based Perturb-seq of alcohol-related genes in SH-SY5Y cells. (A) Proportion of cells successfully perturbed by sgRNA for each sgRNA ID. The 7 genes with any perturbed cells are shown. (B) UMAP dimensional reduction visualization of linear discriminant analysis (LDA) of genes with successfully perturbed cells with NTC cells. (C) Heatmap of gene set enrichment analysis of the change in gene expression in perturbed cells. Heatmap colors indicate the normalized enrichment score (NES) for significantly enriched pathways (adjusted p-value < 0.05). Gene column label colors indicate the direction of association with an alcohol phenotype predicted by MGExA, and pathway row label colors indicate the KEGG pathway category. Heatmap includes KEGG pathways significantly enriched in at least two target genes.

To evaluate the downstream effects of the inhibition of each perturbed gene, we performed gene set enrichment analysis (GSEA) on the differentially expressed genes using pathways from Kyoto Encyclopedia of Genes and Genomes (KEGG)^42^. Of the 186 KEGG pathways used for GSEA, 31 were significantly enriched (FDR < 0.05) by at least one perturbed gene (Table S8), and 16 pathways were enriched by at least two perturbed genes (Figure 4C). Alzheimer’s, Huntington’s, and Parkinson’s disease-related pathways were enriched by multiple CRISPRi target genes. Oxidative phosphorylation was enriched in 5 target genes; oxidative stress may be involved in response to alcohol consumption and AUD^43,44^. The Perturb-seq analysis in the SH-SY5Y cell system suggests that these genes, implicated in alcohol consumption and alcohol use by MPRA, are involved in multiple pathways including important signaling cascades and brain-related diseases.

### Stratification of brain tissue samples by MPRA-derived 3’-UTR activity identified neuroinflammation as a key pathway

To further understand the identified downstream genes and pathways at the tissue level, we utilized genotype and transcriptomics data of pre-frontal cortex from 991 subjects from the CommonMind Consortium (CMC)^45^. For each CommonMind sample, we inferred the 3’-UTR activity of each gene based on the genotypes of 3’-UTR variants and MPRA-derived activity (see Methods). For each gene, we compared global gene expression between samples with the lowest 1/3 and highest 1/3 of these inferred activities, using a generalized linear model with sex and the sample’s institution of origin as covariates. In total, 1,957 (SH-SY5Y) and 8,401 (microglia) unique genes were differentially expressed (FDR < 0.05) between the samples with high and low activities in at least one candidate 3’-UTR identified by MGExA (Table S9). Next, we conducted GSEA of these differentially expressed genes to identify enriched pathways in KEGG (Figure 5). Among the most commonly enriched pathways were Oxidative Phosphorylation, Ribosome, and Parkinson’s disease (Table S10 & S11). Others included toll-like receptor signaling and cytokine-cytokine receptor interaction, two pathways involved in inflammation. Taken together, the cell- and tissue-level analyses of downstream effects suggest that genes identified by combining GWAS and MPRA are also involved in several brain disorders.

**Figure 5.**
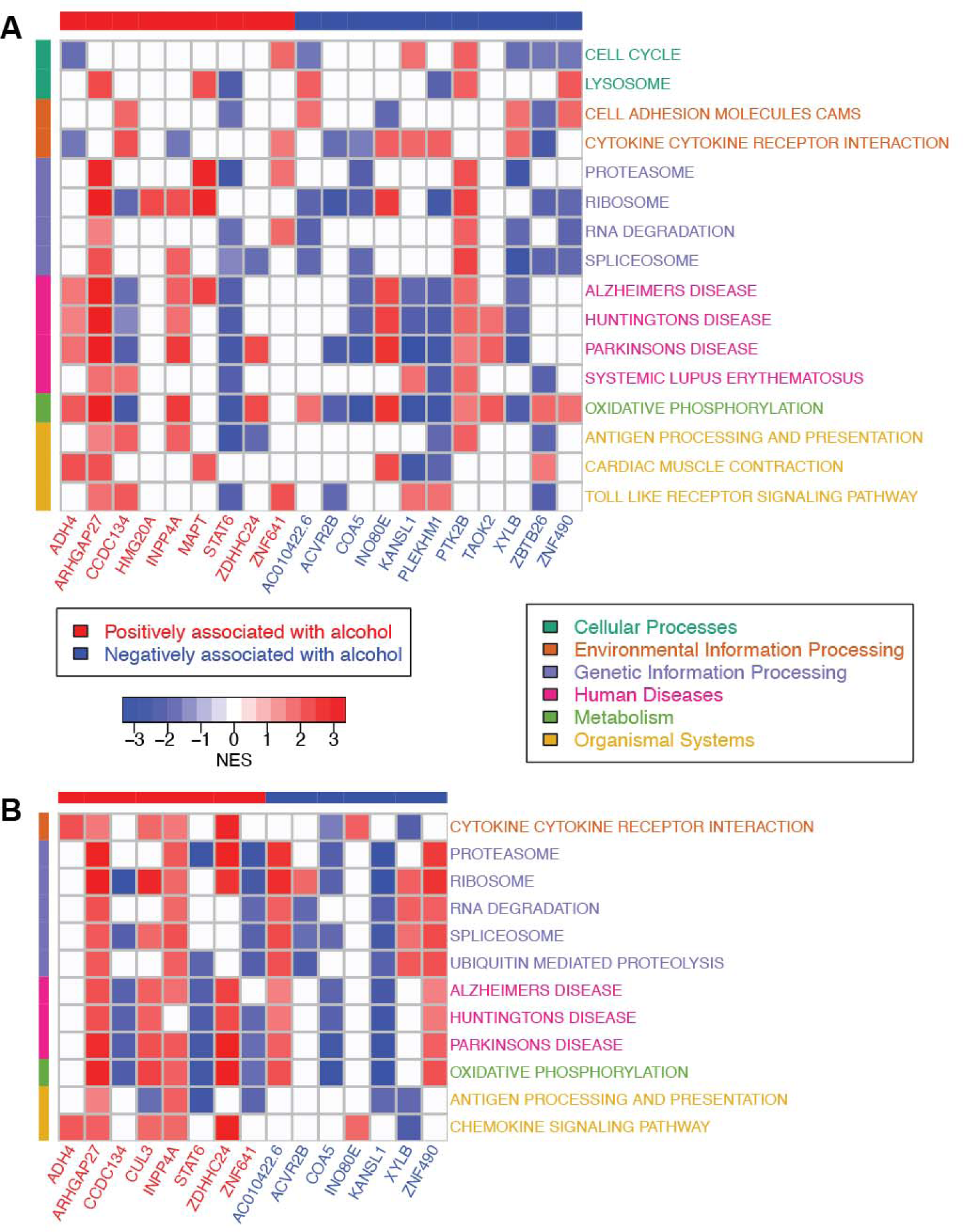
Pathway analysis of differentially expressed genes downstream of the 3’-UTRs of MGExA-identified genes. Heatmaps of the normalized enrichment score (NES) for KEGG pathways that are significantly enriched (adjusted p < 0.05) in the sets of differentially expressed genes downstream of gene 3’-UTRs identified by MGExA to be associated with alcohol consumption or alcohol use disorder. Downstream genes were derived by evaluating differential gene expression in brain tissues stratified by the aggregated effect of the variants in each gene’s 3’-UTR using MPRA variant effects derived from (A) SH-SY5Y and (B) microglia cells. Heatmaps include pathways in KEGG with adjusted p < 0.05 in at least 7 genes and MGExA genes that produced at least 50 downstream differentially expressed genes, though all results are shown in Table S10 and S11. Gene column label colors indicate the direction of association with an alcohol phenotype predicted by MGExA, and pathway row label colors indicate the KEGG pathway category.

In summary, we screened the functional effects of 13,515 3’-UTR variants on gene expression in a neuroblastoma cell line and a microglial cell line and found many that differentially affect gene expression. These functional SNPs explain more of the heritability of alcohol consumption and alcohol use disorder than equivalent random sets selected from the pool of candidate variants we evaluated. We developed a model, MGExA, to aggregate SNP MPRA-derived functional activity with their GWAS effect size to identify genes whose 3’-UTR may contribute to their propensity to increase alcohol consumption or develop AUD. Knocking down these genes in neuronal cells showed that they are involved in oxidative stress pathways, a pathway involved in alcohol consumption and AUD, and other brain disease pathways. By examining differences in tissue-level gene expression between brain samples with different levels of inferred effect of 3’-UTR variants, we found a potential link between neuroinflammation and alcohol use disorder.

## Methods

### Identification of candidate 3’-UTR SNPs

Candidate SNPs were identified from the GWAS catalog^26^ (v1.0, release 2019-08-24, accessed 2019-08-27) and a meta-analysis of alcohol phenotypes^27^ by first selecting lead SNPs that were marginally associated (p < 10^−5^) with traits that mapped to the Experimental Factor Ontology^46^ category “neurological disorders.” For each lead SNP, a linkage disequilibrium (LD) region was defined as the genomic region between the most upstream and downstream distal SNPs with r^2^ > 0.8 within the European populations of the 1000 Genomes Project^28^. A total of 13,515 candidate SNPs were selected from within these LD regions by identifying SNPs within 3’-UTRs defined by dbSNP^47^ build 151 that had minor allele frequency larger than 5% in at least one 1000 Genomes super population.

### Massively Parallel Reporter Assay for 3’-UTR variants

PASSPORT-seq^21,22^, a massively parallel reporter assay (MPRA), was used to evaluate the effect of 3’-UTR SNPs on the expression of a luciferase reporter gene contained in the pIS-0 vector^48^. This method is similar to MPRAu^49^ another MPRA for 3’-UTRs. The overall flow of the protocol is shown in Figure S3. A pool of 24,780 oligonucleotides was synthesized by Agilent (Santa Clara, CA, USA), with each oligonucleotide containing either the reference or alternative allele of 13,515 SNPs flanked by the 25 nucleotides upstream and downstream of its genomic position plus vector-specific sequence at both termini (5’ end: 5’-GCCGTGTAATTCTAGGAGCTC; 3’ end: CGTTCTAGAGTCGGGGCGG-3’) to allow assembly into the pIS-0 plasmid. The oligo pool (10 fmol) was amplified by the polymerase chain reaction (PCR) for 15 cycles with Invitrogen Platinum SuperFi DNA Polymerase mastermix (Thermo Fisher Scientific, Waltham, MA) using 0.25 µM each of PCR primers HJ7207 (5’-GCCGTGTAATTCTAGGAGCTC-3’) and HJ7208 (5’-GCCCCGACTCTAGAACG-3’) in a 50 µL reaction. The products of four independent PCRs were combined and purified using MinElute columns (Qiagen, Germantown, MD).

To construct the plasmid library, the pIS-0 vector^48^ (RRID: Addgene_12178, Addgene, Cambridge, MA) was first linearized using *Sac*I-HF and *Bmt*I-HF restriction enzymes (New England Biolabs (NEB), Ipswich, MA), separated in a 0.8% agarose gel, excised, and purified using a QIAquick gel extraction kit (Qiagen). The linearized vector was assembled with the amplified oligo pool using the NEBuilder HiFi DNA assembly kit (NEB) by mixing 50 ng of the linearized pIS-0 vector and 2 µL of the amplified oligonucleotide pool in a 20 µL reaction volume (Figure S2A). Four independent assembly reactions were conducted, and 3-4 independent transformations were carried out from each assembly reaction, using 1.5 ul of each PCR. Transformation was performed into 50 µL of chemically competent NEB 5-alpha competent *E. coli* cells (NEB), which were spread on two 150-mm LB-agar plates containing 100 µg/ml ampicillin and incubated overnight at 37°C. Bacteria were harvested by adding 4 mL LB to each dish and scraping with an L-shaped cell spreader (Thermo Fisher Scientific). Plasmid DNA was isolated from the bacteria using PureLink HiPure Plasmid Filter Miniprep Kit (Thermo Fisher Scientific) and pooled to produce the final plasmid library.

The MPRA was performed in two cell lines. SH-SY5Y neuroblastoma cells (CRl-2266, ATCC, Manassas, VA) were cultured in a 1:1 mixture of EMEM (American Type Culture Collection; ATCC) and F12K medium (10025-CV, Thermo Fisher Scientific) with 10% (vol/vol) fetal bovine serum (ATCC) and 1% penicillin and streptomycin. Cells were plated in six 10-cm dishes at a density of 5 x 10^6^ cells per dish. After 24 h incubation at 37°C and 5% CO_2_, cells were transfected with X-tremeGENE HP DNA transfection reagent (Millipore Sigma, St. Louis, MO) using 10 µg of plasmid library and 40 µL reagent in 1000 µL Opti-MEM (Thermo Fisher Scientific). A second cell line, SV40-immortalized human microglia cells (Cat.No: T0251, Applied Biological Materials Inc, Richmond, QC, Canada), was cultured in Prigrow III medium with 10% (vol/vol) fetal bovine serum (16000-044, Thermo Fisher Scientific) and 1% penicillin-streptomycin in type I collagen-coated (1:30; Sigma-Aldrich, C3867) six 6-well plates at a density of 4 x 10^5^ cells per wells (Corning™ 3516, Corning, NY). After 24 h, cells were transfected with 1 µg of plasmid library using 3 µl Lipofectamine 3000 reagent (L3000008, Invitrogen, Carlsbad, CA; 3:1 reagent-to-DNA ratio) in Opti-MEM reduced serum medium (31985-070, Thermo Fisher Scientific).

Six independent transfections of each cell line were conducted. After 42 h, the cells were harvested, and DNA and RNA were isolated from each transfection, using the AllPrep DNA/RNA mini kit (Qiagen). Poly-A RNA was isolated using Dynabeads oligo(dT)25 (Thermo Fisher Scientific) and mRNA treated with gDNA wipeout buffer (Qiagen) before preparing sequencing libraries. Sequencing libraries were prepared as described previously^22^ with some modifications. Poly(A)RNA (500 ng) from each independent transfection was reverse transcribed to cDNA using QuantiTech Reverse Transcription kit (Qiagen) using a reporter-specific first strand primer (HJ7211, Table S6 contains primer sequences) that contains the R1 Illumina adapter sequence, a 9-12 nt barcode (with staggered starts to reduce sequencing errors caused by low library complexity) and a 10 nt unique molecular index (UMI, a random sequence to reduce PCR bias). The resulting cDNA was then PCR-amplified with a primer containing the R2 Illumina adapter sequence and one of 6 barcodes, one for each independent transfection (HJ7214-HJ7219, Table S6). Similarly, the DNA was PCR-amplified with another first strand primer (HJ7212) and one of 6 barcodes (HJ7214-HJ7219). The barcoding scheme is shown in Figure S2B. Both RNA and DNA samples underwent a second PCR using primers HJ7220 and HJ7221, after which the resulting libraries were pooled and prepared for sequencing on an Illumina NovaSeq SP (v 1.5) flow cell lane for 121 nt single-end reads.

### Bioinformatics processing and statistical analysis for MPRA data

The pooled samples were differentiated using the barcodes. The FASTQ files were demultiplexed using cutadapt^50^ (v2.7), and the UMI for each read was appended to the read name using umi_tools^51^ (v1.0.0). After trimming the barcodes, UMI, and primer sequences, the remaining oligonucleotide insert sequences (51 nt) were mapped to the reference and alternative alleles of the SNPs. Finally, the reads containing mismatches or duplicated UMIs were discarded (20% of the reads), and the number of unique reads for each sequence was counted using umi_tools.

To calculate the amount of RNA expressed relative to its corresponding plasmid DNA for each oligonucleotide sequence, a generalized linear model implemented within the edgeR package^52,53^ (v. 3.26.8) in R (v. 3.6.0) was used to estimate the coefficient associated with a sequence’s cDNA count compared to what is expected based on its plasmid DNA count (analogous to the ratio of cDNA counts to plasmid DNA counts). The “glmFit” function was used to calculate the coefficients, and the “glmLRT” function was used to perform likelihood ratio tests.

The goal of this MPRA is to evaluate whether the variant allele of a SNP alters the activity of a sequence in the 3’-UTR. A generalized linear mixed model was used:

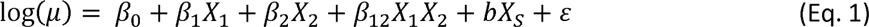

where *µ* is the number of unique reads associated with allele *X*_1_ (reference or alternative), sample type *X*_2_ (DNA or cDNA), and replicate number *X_S_* (1 to 6). The coefficients of the fixed effects are *β*_0_, *β*_1_, *β*_2_, and *β*_12_, while the coefficient for the random effect is *b*. The effect of the variant on gene expression was estimated using *β*_12_, the coefficient of the interaction between allele and sample type. Our null hypothesis (*H*_0_) is *β*_12_ =0, analogous to stating that the relative counts of the reference and alternative alleles in the cDNA is equal to that in the plasmid DNA. A false discovery rate (FDR; Benjamini-Hochberg procedure^54^) < 0.05 was used as the threshold for significance.

### Heritability enrichment analysis of MPRA-derived significant SNPs

Heritability enrichment analysis was conducted using LDAK^55^ (v. 5.0) by comparing the heritability contribution of SNPs significant in our MPRA (FDR ≤ 0.05; ref. ^54^) with the heritability contribution of 1000 sets containing the same number of candidate SNPs randomly drawn from the full 13,515 SNPs. LDAK estimates the expected heritability contribution of each GWAS SNP using the GWAS summary statistics, linkage disequilibrium patterns, and minor allele frequency for each SNP. A permutation p-value was determined by calculating the fraction of the 1000 values that were greater than the share of heritability of the FDR-significant SNPs.

### MPRA-mediated Gene Expression Association Analysis (MGExA) to identify genes whose 3’-UTRs are associated with Alcohol Consumption and AUD

We developed a computational framework, MPRA-mediated Gene Expression Association Analysis (MGExA), to evaluate the association of the 3’-UTR component of gene expression with a phenotype. First, we defined for each gene in an individual the genetic component of gene expression due to 3’-UTR effects by summing the MPRA-derived effects of the 3’-UTR SNPs weighted by alternative allele counts of the SNPs:

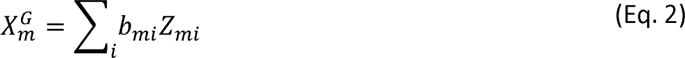

where 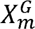 is the calculated genetic component of 3’-UTR activity for gene *m*, *b_mi_* is the MPRA-derived effect of SNP *i*, and *Z_mi_* is the number of alternative alleles of SNP *i*. Here, *b_mi_* is defined as the z value associated with *β*_12_ from Eq. 1. A GWAS-like approach could be used to determine whether the genetic component of a gene’s 3’-GWAS activity is associated with a trait of interest. However, this requires individual-level data for each target phenotype. Inspired by Summary Mendelian Randomization^25^, Transcriptome-Wide Association Study (TWAS)^24^, and S-Predixcan^56^, our approach instead utilizes GWAS summary statistics for a disease of interest and uses individual-level data from a reference population to estimate the variance of 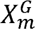 in the GWAS population. Here we used the 2504 samples from phase 3 of the 1000 Genomes Project^28^ as the reference population. To evaluate the association of the 3’-UTR component of gene expression with a phenotype using GWAS summary statistics, we calculated the following test statistic:

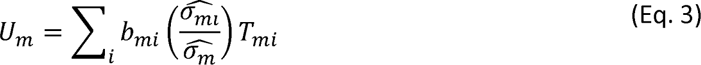

where *U_m_* is the Z-score for gene *m*, *b_mi_* is the MPRA-derived effect of SNP i, 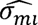 is the standard deviation of the alternative allele count *Z_mi_* of SNP *i* for individuals in the reference population, 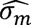 is the standard deviation of 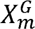 (the individual-level genetic component of 3’-UTR activity, Eq. 2) for all individuals in the reference population, and *T_mi_* is the test statistic derived from GWAS summary statistics by dividing the effect size by the standard error. A p-value was derived by assuming *U_m_* to be normally distributed.

To apply MGExA to a GWAS, we collected the SNPs evaluated in both the MPRA and the GWAS. For each gene containing at least one SNP, we calculated *U_m_* We applied MGExA to genes with SNPs evaluated with our MRPA and GWAS summary statistics of alcoholic drinks per week from GSCAN^6^ The significant genes (FDR < 0.2) were then replicated with a GWAS of a broader alcohol consumption phenotype, AUDIT-C, from the Million Veteran Program (MVP)^2^. Similarly, we identified genes associated with alcohol use disorder by initially applying MGExA to GWAS summary statistics of AUD (defined by ICD codes) from MVP^2^, and further evaluated the significant genes (FDR < 0.2) using UK BioBank (UKBB) AUDIT-P (AUDIT-Problems)^1^.

### Generation of an SH-SY5Y cell line that stably expresses KRAB-dCas9

SH-SY5Y cells stably expressing KRAB-dCas9 were generated following the protocol for CRISPRi by Sigma-Aldrich. Briefly, cells were seeded in 6-well culture plates at 2 x 10^5^ cells per well (10% confluency) and incubated at 37°C for 24 h. The medium was replaced with growth medium containing 8 µg/mL polybrene, cells were transduced with a KRAB-dCas9 lentiviral construct at multiplicity of infection (MOI) 0.2. After 24 h, media was replaced with fresh growth medium, and the cells were incubated for another 24 h. To begin antibiotic selection, medium was replaced with fresh medium containing Blasticidin (5 µg/mL). Antibiotic-containing medium was replaced every other day for 8 days, and the resulting cell line was cryopreserved for future experiments.

### Perturb-seq single cell CRISPR-induced inhibition of target genes

We performed Perturb-seq^40^ with candidate genes derived from MGExA. For each of these 31 genes, 5 guide RNA sequences were generated to target the region near the transcriptional start site of each gene. Additionally, 18 positive controls and 18 negative controls were also included, for a total of 191 sgRNAs in the CRISPRi library. The pooled lentiviral construct for the library was generated by Sigma-Aldrich.

Using the KRAB-dCas9 SH-SY5Y cell line, cells were seeded at 3 x 10^6^ cells per dish. After 24 h incubation, cells were transduced with the pooled lentiviral construct at a target MOI of 0.2. After another 24 h, medium was changed, then after 24 h incubation, antibiotic selection began using medium containing 1.5 µg/mL Puromycin. Medium with antibiotic was changed every other day for 7 more days. Cells were then harvested for single cell sequencing library preparation using 10X Genomics Chromium X.

Approximately 33,000 cells were loaded to target 20,000 cells in each well of a chip N, and two wells were used. The libraries were constructed following the 10X Genomics protocol for Chromium NEXT GEM Single Cell 5’ HT Reagent Kits (Dual Index) with Feature Barcode technology for CRISPR Screening. To identify the sgRNAs present in each gel bead, CRISPR primers were included in the reverse transcription step to amplify sgRNA as well as mRNA. After cDNA amplification, cDNA from sgRNA was separated from mRNA using size selection with SPRISelect and prepared for sequencing. The final cDNA and CRIPSR libraries were sequenced on an Illumina NovaSeq 6000. The cellranger multi algorithm of Cell Ranger 7.0 (http://support.10xgenomics.com/) was used to process the sequence data. Of the 30,972 cell barcodes identified, 18,507 contained at least one sgRNA barcode. Of these, 8,842 cells were identified to have been transduced with one target gene sgRNA, and 1,331 cells were identified to be transduced with NTC.

To identify which cells were successfully perturbed, we used Seurat’s mixscape function^41^ closely following their recommended vignette. Briefly, the sequencing results for cells with only one sgRNA were loaded into Seurat and normalized using SCTransform^57^. The perturbation signature of each cell was calculated using the CalcPerturbSig function using ndims = 40 and num.neighbors=20 with “pca” as the dimensional reduction method. Only 7 genes were successfully perturbed. The RunMixscape function was then run with default parameters using default pooling mode then fine mode. Default mode treats all five sgRNA IDs for one target gene as the same class, while fine mode treats each sgRNA ID as its own class. To evaluate differential expression for each target gene, the expression of perturbed cells was compared to NTC cells using Wilcoxon Rank-Sum test with the FindMarkers function. Perturbation responses across cells were visualized using Linear Discriminant Analysis (LDA), a method that maximizes the separation between labels. The MixscapeLDA function was used to do dimensional reduction, and RunUMAP was used to visualize the resulting UMAP clusters.

### Identifying relevant pathways using gene set enrichment analysis

We performed gene set enrichment analysis (GSEA) using the fgsea^58^ package in R for each set of expression changes from either target gene perturbation or MPRA-derived gene activity stratification using pathways from Kyoto Encyclopedia of Genes and Genomes (KEGG)^42^. For each analysis, we used a list of genes ordered by −log10(p-value of differential expression analysis) multiplied by the direction of effect as the gene set input. We applied Benjamini-Hochberg correction to report the false discovery rate of each pathway.

### Differential Expression and Pathway Analysis of Genes Downstream of MPRA-derived 3’-UTR Activity

RNA sequencing data from 991 brain samples were obtained from the CommonMind Consortium^45^ (accessed 7/6/2020). For each MPRA-evaluated SNP in each brain sample, genotypes were determined from the RNA-seq data by calculating the frequency of each allele, where loci with the minor allele being less than 5% were considered homozygous and all else were heterozygous. Using Eq. 2, the genetic component of 3’-UTR activity for each gene of interest was determined for each sample. For each gene of interest, the activity score range was stratified into thirds to identify groups of low and high 3’-UTR activity, and a generalize linear model using the glmFit^53^ function of edgeR^52^ (v. 3.14) was used to perform gene expression analysis comparing these two groups, using sex and the sample’s institution of origin as covariates. For each gene of interest identified by MGExA, a set of differentially expressed genes was constructed, and over-represented KEGG pathways^42^ in each gene set were identified using g:Profiler^59^ (using FDR < 0.05 as the significance threshold).

## Author Contributions

Conceptualization: ABC, HJE, YL; Methodology: ABC, XY, KST, HG, JLR, XX, YW, HJE, YL; Software: ABC, HG, DL; Investigation: ABC, XY, KST, APT; Writing – Original Draft: ABC, XY, HJE, YL; Writing – Review & Editing: ABC, HJE, YL; Supervision: GEL, HG, YW, TMF, JAT, HJE, YL; Funding Acquisition: TMF, JAT, HJE, YL.

## Acknowledgements

This study is supported by the National Institute on Drug Abuse grant R01DA053722 (to HJE and YL), National Institute on Alcohol Abuse and Alcoholism grant U10AA08403 (COGA), and the National Center for Advancing Translational Sciences, Clinical and Translational Sciences Award Grant TL1TR002531 (T. Hurley, PI). High-throughput sequencing was carried out in the Center for Medical Genomics at Indiana University School of Medicine, which is partially supported by the Indiana University Grand Challenges Precision Health Initiative.

Bio-samples and/or data for this publication were obtained from NIMH Repository & Genomics Resource, a centralized national biorepository for genetic studies of psychiatric disorders. Data were generated as part of the CommonMind Consortium supported by funding from Takeda Pharmaceuticals Company Limited, F. Hoffman-La Roche Ltd and NIH grants R01MH085542, R01MH093725, P50MH066392, P50MH080405, R01MH097276, RO1-MH-075916, P50M096891, P50MH084053S1, R37MH057881, AG02219, AG05138, MH06692, R01MH110921, R01MH109677, R01MH109897, U01MH103392, and contract HHSN271201300031C through IRP NIMH. Brain tissue for the study was obtained from the following brain bank collections: the Mount Sinai NIH Brain and Tissue Repository, the University of Pennsylvania Alzheimer’s Disease Core Center, the University of Pittsburgh NeuroBioBank and Brain and Tissue Repositories, and the NIMH Human Brain Collection Core. CMC Leadership: Panos Roussos, Joseph Buxbaum, Andrew Chess, Schahram Akbarian, Vahram Haroutunian (Icahn School of Medicine at Mount Sinai), Bernie Devlin, David Lewis (University of Pittsburgh), Raquel Gur, Chang-Gyu Hahn (University of Pennsylvania), Enrico Domenici (University of Trento), Mette A. Peters, Solveig Sieberts (Sage Bionetworks), Thomas Lehner, Stefano Marenco, Barbara K. Lipska (NIMH).

## Data and code availability

The datasets and code generated during this study will be available upon publication.

## References

1. Sanchez-Roige, S., Palmer, A.A., Fontanillas, P., Elson, S.L., Adams, M.J., Howard, D.M., Edenberg, H.J., Davies, G., Crist, R.C., Deary, I.J., et al. (2019). Genome-Wide Association Study Meta-Analysis of the Alcohol Use Disorders Identification Test (AUDIT) in Two Population-Based Cohorts. Am J Psychiatry 176, 107–118. 10.1176/appi.ajp.2018.18040369.

2. Kranzler, H.R., Zhou, H., Kember, R.L., Vickers Smith, R., Justice, A.C., Damrauer, S., Tsao, P.S., Klarin, D., Baras, A., Reid, J., et al. (2019). Genome-wide association study of alcohol consumption and use disorder in 274,424 individuals from multiple populations. Nature Communications 10, 1499. 10.1038/s41467-019-09480-8.

3. Gelernter, J., Kranzler, H.R., Sherva, R., Almasy, L., Koesterer, R., Smith, A.H., Anton, R., Preuss, U.W., Ridinger, M., Rujescu, D., et al. (2014). Genome-wide association study of alcohol dependence:significant findings in African- and European-Americans including novel risk loci. Mol Psychiatry 19, 41–49. 10.1038/mp.2013.145.

4. Zhou, H., Kalayasiri, R., Sun, Y., Nunez, Y.Z., Deng, H.W., Chen, X.D., Justice, A.C., Kranzler, H.R., Chang, S., Lu, L., et al. (2022). Genome-wide meta-analysis of alcohol use disorder in East Asians. Neuropsychopharmacology 47, 1791–1797. 10.1038/s41386-022-01265-w.

5. Clarke, T.K., Adams, M.J., Davies, G., Howard, D.M., Hall, L.S., Padmanabhan, S., Murray, A.D., Smith, B.H., Campbell, A., Hayward, C., et al. (2017). Genome-wide association study of alcohol consumption and genetic overlap with other health-related traits in UK Biobank (N=1121117). Mol Psychiatry 22, 1376–1384. 10.1038/mp.2017.153.

6. Liu, M., Jiang, Y., Wedow, R., Li, Y., Brazel, D.M., Chen, F., Datta, G., Davila-Velderrain, J., McGuire, D., Tian, C., et al. (2019). Association studies of up to 1.2 million individuals yield new insights into the genetic etiology of tobacco and alcohol use. Nature Genetics 51, 237–244. 10.1038/s41588-018-0307-5.

7. Schumann, G., Liu, C., O’Reilly, P., Gao, H., Song, P., Xu, B., Ruggeri, B., Amin, N., Jia, T., Preis, S., et al. (2016). KLB is associated with alcohol drinking, and its gene product β-Klotho is necessary for FGF21 regulation of alcohol preference. Proc Natl Acad Sci U S A 113, 14372–14377. 10.1073/pnas.1611243113.

8. Jorgenson, E., Thai, K.K., Hoffmann, T.J., Sakoda, L.C., Kvale, M.N., Banda, Y., Schaefer, C., Risch, N., Mertens, J., Weisner, C., and Choquet, H. (2017). Genetic contributors to variation in alcohol consumption vary by race/ethnicity in a large multi-ethnic genome-wide association study. Molecular Psychiatry 22, 1359–1367. 10.1038/mp.2017.101.

9. Mallard, T.T., Savage, J.E., Johnson, E.C., Huang, Y., Edwards, A.C., Hottenga, J.J., Grotzinger, A.D., Gustavson, D.E., Jennings, M.V., Anokhin, A., et al. (2022). Item-Level Genome-Wide Association Study of the Alcohol Use Disorders Identification Test in Three Population-Based Cohorts. Am J Psychiatry 179, 58–70. 10.1176/appi.ajp.2020.20091390.

10. Lai, D., Kapoor, M., Wetherill, L., Schwandt, M., Ramchandani, V.A., Goldman, D., Chao, M., Almasy, L., Bucholz, K., Hart, R.P., et al. (2021). Genome-wide admixture mapping of DSM-IV alcohol dependence, criterion count, and the self-rating of the effects of ethanol in African American populations. Am J Med Genet B Neuropsychiatr Genet 186, 151–161. 10.1002/ajmg.b.32805.

11. Walters, R.K., Polimanti, R., Johnson, E.C., McClintick, J.N., Adams, M.J., Adkins, A.E., Aliev, F., Bacanu, S.A., Batzler, A., Bertelsen, S., et al. (2018). Transancestral GWAS of alcohol dependence reveals common genetic underpinnings with psychiatric disorders. Nat Neurosci 21, 1656–1669. 10.1038/s41593-018-0275-1.

12. Wetherill, L., Lai, D., Johnson, E.C., Anokhin, A., Bauer, L., Bucholz, K.K., Dick, D.M., Hariri, A.R., Hesselbrock, V., Kamarajan, C., et al. (2019). Genome-wide association study identifies loci associated with liability to alcohol and drug dependence that is associated with variability in reward-related ventral striatum activity in African- and European-Americans. Genes Brain Behav 18, e12580. 10.1111/gbb.12580.

13. Kember, R.L., Vickers-Smith, R., Zhou, H., Xu, H., Jennings, M., Dao, C., Davis, L., Sanchez-Roige, S., Justice, A.C., Gelernter, J., et al. (2023). Genetic Underpinnings of the Transition From Alcohol Consumption to Alcohol Use Disorder: Shared and Unique Genetic Architectures in a Cross-Ancestry Sample. Am J Psychiatry 180, 584–593. 10.1176/appi.ajp.21090892.

14. Lai, D., Wetherill, L., Kapoor, M., Johnson, E.C., Schwandt, M., Ramchandani, V.A., Goldman, D., Joslyn, G., Rao, X., Liu, Y., et al. (2020). Genome-wide association studies of the self-rating of effects of ethanol (SRE). Addict Biol 25, e12800. 10.1111/adb.12800.

15. Zhou, H., Kember, R.L., Deak, J.D., Xu, H., Toikumo, S., Yuan, K., Lind, P.A., Farajzadeh, L., Wang, L., Hatoum, A.S., et al. (2023). Multi-ancestry study of the genetics of problematic alcohol use in >1 million individuals. medRxiv. 10.1101/2023.01.24.23284960.

16. Zhou, H., Sealock, J.M., Sanchez-Roige, S., Clarke, T.K., Levey, D.F., Cheng, Z., Li, B., Polimanti, R., Kember, R.L., Smith, R.V., et al. (2020). Genome-wide meta-analysis of problematic alcohol use in 435,563 individuals yields insights into biology and relationships with other traits. Nat Neurosci 23, 809–818. 10.1038/s41593-020-0643-5.

17. Barr, P.B., Driver, M.N., Kuo, S.I., Stephenson, M., Aliev, F., Linner, R.K., Marks, J., Anokhin, A.P., Bucholz, K., Chan, G., et al. (2022). Clinical, environmental, and genetic risk factors for substance use disorders: characterizing combined effects across multiple cohorts. Mol Psychiatry 27, 4633–4641. 10.1038/s41380-022-01801-6.

18. Hatoum, A.S., Colbert, S.M.C., Johnson, E.C., Huggett, S.B., Deak, J.D., Pathak, G., Jennings, M.V., Paul, S.E., Karcher, N.R., Hansen, I., et al. (2023). Multivariate genome-wide association meta-analysis of over 1 million subjects identifies loci underlying multiple substance use disorders. Nat Ment Health 1, 210–223. 10.1038/s44220-023-00034-y.

19. Melnikov, A., Murugan, A., Zhang, X., Tesileanu, T., Wang, L., Rogov, P., Feizi, S., Gnirke, A., Callan, C.G., Jr., Kinney, J.B., et al. (2012). Systematic dissection and optimization of inducible enhancers in human cells using a massively parallel reporter assay. Nat Biotechnol 30, 271–277. 10.1038/nbt.2137.

20. Kheradpour, P., Ernst, J., Melnikov, A., Rogov, P., Wang, L., Zhang, X., Alston, J., Mikkelsen, T.S., and Kellis, M. (2013). Systematic dissection of regulatory motifs in 2000 predicted human enhancers using a massively parallel reporter assay. Genome Res 23, 800–811. 10.1101/gr.144899.112.

21. Ipe, J., Collins, K.S., Hao, Y., Gao, H., Bhatia, P., Gaedigk, A., Liu, Y., and Skaar, T.C. (2018). PASSPORT-seq: A Novel High-Throughput Bioassay to Functionally Test Polymorphisms in Micro-RNA Target Sites. Front Genet 9, 219. 10.3389/fgene.2018.00219.

22. Rao, X., Thapa, K.S., Chen, A.B., Lin, H., Gao, H., Reiter, J.L., Hargreaves, K.A., Ipe, J., Lai, D., Xuei, X., et al. (2021). Allele-specific expression and high-throughput reporter assay reveal functional genetic variants associated with alcohol use disorders. Mol Psychiatry 26, 1142–1151. 10.1038/s41380-019-0508-z.

23. Gamazon, E.R., Wheeler, H.E., Shah, K.P., Mozaffari, S.V., Aquino-Michaels, K., Carroll, R.J., Eyler, A.E., Denny, J.C., Consortium, G.T., Nicolae, D.L., et al. (2015). A gene-based association method for mapping traits using reference transcriptome data. Nature genetics 47, 1091–1098. 10.1038/ng.3367.

24. Gusev, A., Ko, A., Shi, H., Bhatia, G., Chung, W., Penninx, B.W., Jansen, R., de Geus, E.J., Boomsma, D.I., Wright, F.A., et al. (2016). Integrative approaches for large-scale transcriptome-wide association studies. Nat Genet 48, 245–252. 10.1038/ng.3506.

25. Zhu, Z., Zhang, F., Hu, H., Bakshi, A., Robinson, M.R., Powell, J.E., Montgomery, G.W., Goddard, M.E., Wray, N.R., Visscher, P.M., and Yang, J. (2016). Integration of summary data from GWAS and eQTL studies predicts complex trait gene targets. Nat Genet 48, 481–487. 10.1038/ng.3538.

26. Buniello, A., MacArthur, J.A.L., Cerezo, M., Harris, L.W., Hayhurst, J., Malangone, C., McMahon, A., Morales, J., Mountjoy, E., Sollis, E., et al. (2019). The NHGRI-EBI GWAS Catalog of published genome-wide association studies, targeted arrays and summary statistics 2019. Nucleic Acids Res 47, D1005–d1012. 10.1093/nar/gky1120.

27. Lai, D., Wetherill, L., Bertelsen, S., Carey, C.E., Kamarajan, C., Kapoor, M., Meyers, J.L., Anokhin, A.P., Bennett, D.A., Bucholz, K.K., et al. (2019). Genome-wide association studies of alcohol dependence, DSM-IV criterion count and individual criteria. Genes Brain Behav 18, e12579. 10.1111/gbb.12579.

28. Auton, A., Abecasis, G.R., Altshuler, D.M., Durbin, R.M., Abecasis, G.R., Bentley, D.R., Chakravarti, A., Clark, A.G., Donnelly, P., Eichler, E.E., et al. (2015). A global reference for human genetic variation. Nature 526, 68–74. 10.1038/nature15393.

29. Biedler, J.L., Helson, L., and Spengler, B.A. (1973). Morphology and growth, tumorigenicity, and cytogenetics of human neuroblastoma cells in continuous culture. Cancer Res 33, 2643–2652.

30. Siletti, K., Hodge, R., Mossi Albiach, A., Lee, K.W., Ding, S.L., Hu, L., Lonnerberg, P., Bakken, T., Casper, T., Clark, M., et al. (2023). Transcriptomic diversity of cell types across the adult human brain. Science 382, eadd7046. 10.1126/science.add7046.

31. Luo, X., Kranzler, H.R., Zuo, L., Lappalainen, J., Yang, B.-z., and Gelernter, J. (2006). ADH4 Gene Variation is Associated with Alcohol Dependence and Drug Dependence in European Americans: Results from HWD Tests and Case–Control Association Studies. Neuropsychopharmacology 31, 1085–1095. 10.1038/sj.npp.1300925.

32. Zhong, V.W., Kuang, A., Danning, R.D., Kraft, P., van Dam, R.M., Chasman, D.I., and Cornelis, M.C. (2019). A genome-wide association study of bitter and sweet beverage consumption. Hum Mol Genet 28, 2449–2457. 10.1093/hmg/ddz061.

33. Evangelou, E., Gao, H., Chu, C., Ntritsos, G., Blakeley, P., Butts, A.R., Pazoki, R., Suzuki, H., Koskeridis, F., Yiorkas, A.M., et al. (2019). New alcohol-related genes suggest shared genetic mechanisms with neuropsychiatric disorders. Nat Hum Behav 3, 950–961. 10.1038/s41562-019-0653-z.

34. olde Heuvel, F., Holl, S., Chandrasekar, A., Li, Z., Wang, Y., Rehman, R., Förstner, P., Sinske, D., Palmer, A., Wiesner, D., et al. (2019). STAT6 mediates the effect of ethanol on neuroinflammatory response in TBI. Brain, Behavior, and Immunity 81, 228–246. 10.1016/j.bbi.2019.06.019.

35. Kapfhamer, D., Taylor, S., Zou, M.E., Lim, J.P., Kharazia, V., and Heberlein, U. (2013). Taok2 controls behavioral response to ethanol in mice. Genes, Brain and Behavior 12, 87–97. 10.1111/j.1601-183X.2012.00834.x.

36. Gerring, Z.F., Vargas, A.M., Gamazon, E.R., and Derks, E.M. (2021). An integrative systems-based analysis of substance use: eQTL-informed gene-based tests, gene networks, and biological mechanisms. American Journal of Medical Genetics Part B: Neuropsychiatric Genetics 186, 162–172. 10.1002/ajmg.b.32829.

37. Gaddis, N., Mathur, R., Marks, J., Zhou, L., Quach, B., Waldrop, A., Levran, O., Agrawal, A., Randesi, M., Adelson, M., et al. (2022). Multi-trait genome-wide association study of opioid addiction: OPRM1 and beyond. Sci Rep 12, 16873. 10.1038/s41598-022-21003-y.

38. Lambert, J.-C., Ibrahim-Verbaas, C.A., Harold, D., Naj, A.C., Sims, R., Bellenguez, C., Jun, G., DeStefano, A.L., Bis, J.C., Beecham, G.W., et al. (2013). Meta-analysis of 74,046 individuals identifies 11 new susceptibility loci for Alzheimer’s disease. Nature Genetics 45, 1452–1458. 10.1038/ng.2802.

39. Le Hellard, S., Mühleisen, T.W., Djurovic, S., Fernø, J., Ouriaghi, Z., Mattheisen, M., Vasilescu, C., Raeder, M.B., Hansen, T., Strohmaier, J., et al. (2010). Polymorphisms in SREBF1 and SREBF2, two antipsychotic-activated transcription factors controlling cellular lipogenesis, are associated with schizophrenia in German and Scandinavian samples. Molecular Psychiatry 15, 463–472. 10.1038/mp.2008.110.

40. Dixit, A., Parnas, O., Li, B., Chen, J., Fulco, C.P., Jerby-Arnon, L., Marjanovic, N.D., Dionne, D., Burks, T., Raychowdhury, R., et al. (2016). Perturb-Seq: Dissecting Molecular Circuits with Scalable Single-Cell RNA Profiling of Pooled Genetic Screens. Cell 167, 1853–1866 e1817. 10.1016/j.cell.2016.11.038.

41. Papalexi, E., Mimitou, E.P., Butler, A.W., Foster, S., Bracken, B., Mauck, W.M., 3rd, Wessels, H.H., Hao, Y., Yeung, B.Z., Smibert, P., and Satija, R. (2021). Characterizing the molecular regulation of inhibitory immune checkpoints with multimodal single-cell screens. Nat Genet 53, 322–331. 10.1038/s41588-021-00778-2.

42. Kanehisa, M., Furumichi, M., Sato, Y., Kawashima, M., and Ishiguro-Watanabe, M. (2023). KEGG for taxonomy-based analysis of pathways and genomes. Nucleic Acids Res 51, D587–D592. 10.1093/nar/gkac963.

43. Das, S.K., and Vasudevan, D.M. (2007). Alcohol-induced oxidative stress. Life Sci 81, 177–187. 10.1016/j.lfs.2007.05.005.

44. Kamal, H., Tan, G.C., Ibrahim, S.F., Shaikh, M.F., Mohamed, I.N., Mohamed, R.M.P., Hamid, A.A., Ugusman, A., and Kumar, J. (2020). Alcohol Use Disorder, Neurodegeneration, Alzheimer’s and Parkinson’s Disease: Interplay Between Oxidative Stress, Neuroimmune Response and Excitotoxicity. Front Cell Neurosci 14, 282. 10.3389/fncel.2020.00282.

45. Hoffman, G.E., Bendl, J., Voloudakis, G., Montgomery, K.S., Sloofman, L., Wang, Y.-C., Shah, H.R., Hauberg, M.E., Johnson, J.S., Girdhar, K., et al. (2019). CommonMind Consortium provides transcriptomic and epigenomic data for Schizophrenia and Bipolar Disorder. Scientific Data 6, 180. 10.1038/s41597-019-0183-6.

46. Malone, J., Holloway, E., Adamusiak, T., Kapushesky, M., Zheng, J., Kolesnikov, N., Zhukova, A., Brazma, A., and Parkinson, H. (2010). Modeling sample variables with an Experimental Factor Ontology. Bioinformatics 26, 1112–1118. 10.1093/bioinformatics/btq099.

47. Sherry, S.T., Ward, M.H., Kholodov, M., Baker, J., Phan, L., Smigielski, E.M., and Sirotkin, K. (2001). dbSNP: the NCBI database of genetic variation. Nucleic Acids Res 29, 308–311. 10.1093/nar/29.1.308.

48. Yekta, S., Shih, I.H., and Bartel, D.P. (2004). MicroRNA-directed cleavage of HOXB8 mRNA. Science 304, 594–596. 10.1126/science.1097434.

49. Griesemer, D., Xue, J.R., Reilly, S.K., Ulirsch, J.C., Kukreja, K., Davis, J.R., Kanai, M., Yang, D.K., Butts, J.C., Guney, M.H., et al. (2021). Genome-wide functional screen of 3’UTR variants uncovers causal variants for human disease and evolution. Cell 184, 5247–5260.e5219. 10.1016/j.cell.2021.08.025.

50. Martin, M. (2011). Cutadapt removes adapter sequences from high-throughput sequencing reads. 2011 17, 3. 10.14806/ej.17.1.200.

51. Smith, T., Heger, A., and Sudbery, I. (2017). UMI-tools: modeling sequencing errors in Unique Molecular Identifiers to improve quantification accuracy. Genome Res 27, 491–499. 10.1101/gr.209601.116.

52. Robinson, M.D., McCarthy, D.J., and Smyth, G.K. (2010). edgeR: a Bioconductor package for differential expression analysis of digital gene expression data. Bioinformatics 26, 139–140. 10.1093/bioinformatics/btp616.

53. McCarthy, D.J., Chen, Y., and Smyth, G.K. (2012). Differential expression analysis of multifactor RNA-Seq experiments with respect to biological variation. Nucleic Acids Res 40, 4288–4297. 10.1093/nar/gks042.

54. Benjamini, Y., and Hochberg, Y. (1995). Controlling the False Discovery Rate: A Practical and Powerful Approach to Multiple Testing. Journal of the Royal Statistical Society: Series B (Methodological) 57, 289–300. 10.1111/j.2517-6161.1995.tb02031.x.

55. Speed, D., Holmes, J., and Balding, D.J. (2020). Evaluating and improving heritability models using summary statistics. Nature Genetics 52, 458–462. 10.1038/s41588-020-0600-y.

56. Barbeira, A.N., Dickinson, S.P., Bonazzola, R., Zheng, J., Wheeler, H.E., Torres, J.M., Torstenson, E.S., Shah, K.P., Garcia, T., Edwards, T.L., et al. (2018). Exploring the phenotypic consequences of tissue specific gene expression variation inferred from GWAS summary statistics. Nature Communications 9, 1825. 10.1038/s41467-018-03621-1.

57. Choudhary, S., and Satija, R. (2022). Comparison and evaluation of statistical error models for scRNA-seq. Genome Biol 23, 27. 10.1186/s13059-021-02584-9.

58. Korotkevich, G., Sukhov, V., Budin, N., Shpak, B., Artyomov, M.N., and Sergushichev, A. (2021). Fast gene set enrichment analysis. bioRxiv, 060012. 10.1101/060012.

59. Reimand, J., Kull, M., Peterson, H., Hansen, J., and Vilo, J. (2007). g:Profiler--a web-based toolset for functional profiling of gene lists from large-scale experiments. Nucleic Acids Res 35, W193–200. 10.1093/nar/gkm226.

