## Supplemental Information for "Functional 3’-UTR Variants Identify Regulatory Mechanisms Impacting Alcohol Use Disorder and Related Traits"

**Supplemental Figures**

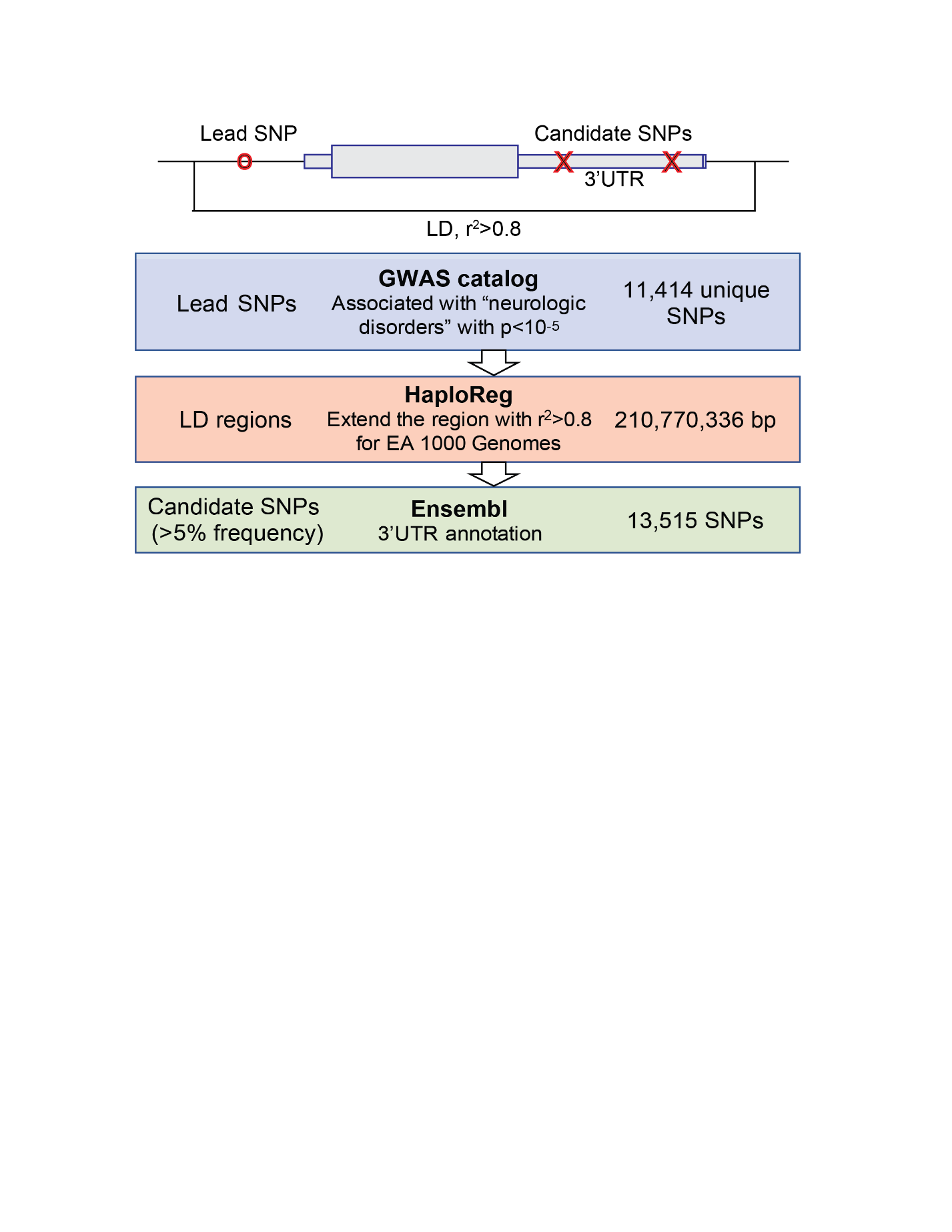

**Figure S1. Overview of the variant selection strategy.**

First, lead SNPs were chosen for marginal association (p < 10^-5^) with neurological disorders^1^ and AUD^2^. Second, for each SNP, an LD region was defined as the region between the furthest two SNPs that had r^2^>0.8 with the lead SNP in the European-American population of 1000 Genomes [cite]. Finally, 3’UTR SNPs with minor allele frequencies larger than 5% in at least one 1000 Genomes super-population were identified based on annotations in dbSNP^3^ build 151 for evaluation by the MPRA.

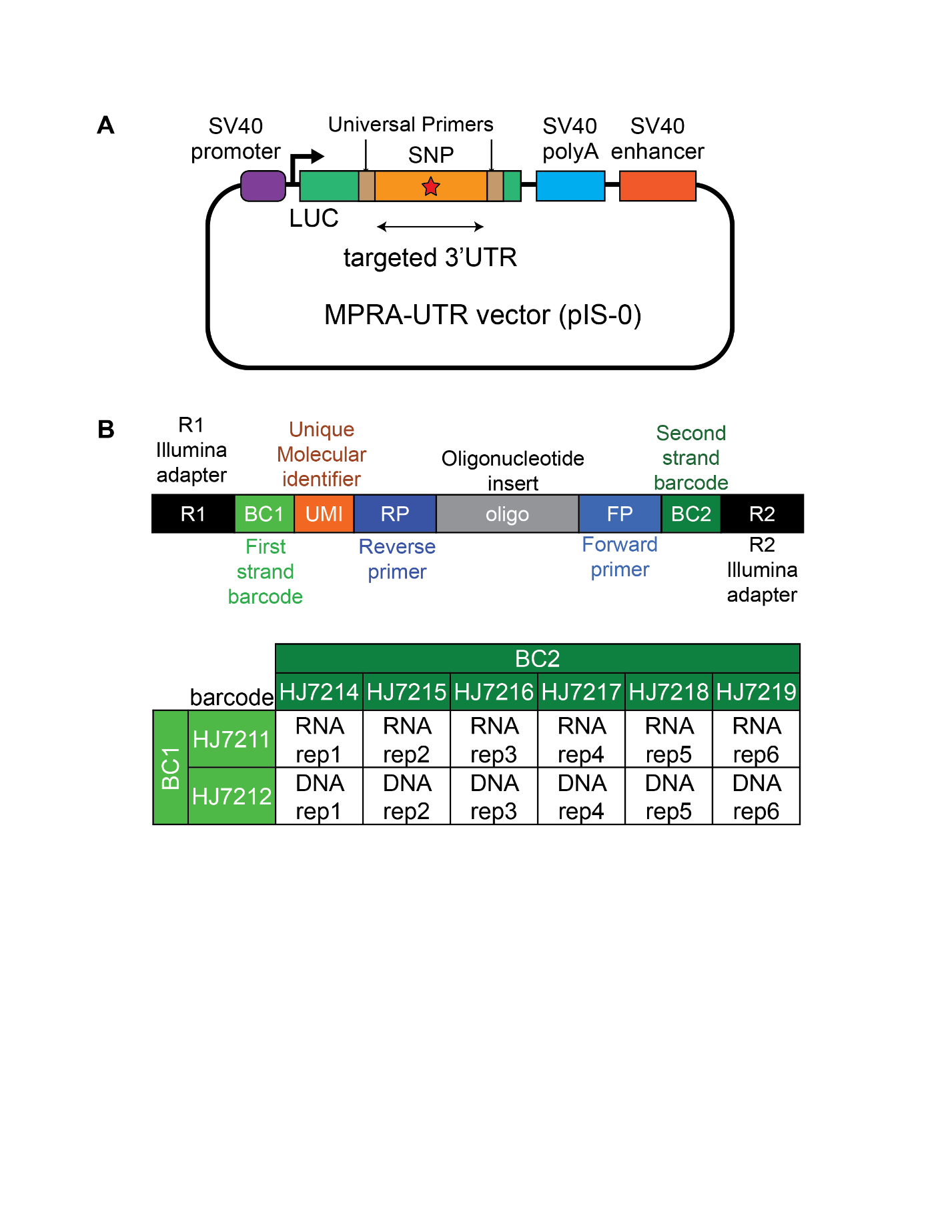

**Figure S2. Schematic summary of the MPRA plasmid and sequencing strategy.** (a) Diagram of the pIS-0 vector used in MPRA. The sequence of interest is inserted into the 3’-UTR of the luciferase (LUC) gene flanked by universal primers that allow for plasmid assembly. (b) A schematic of the assembly strategy for sequencing. Each target nucleotide sequence isolated from RNA and DNA is barcoded to label its nucleic acid type and sample replicate number. The barcoding combinations are shown in the table. Unique molecular identifiers are added to eliminate PCR amplification bias.

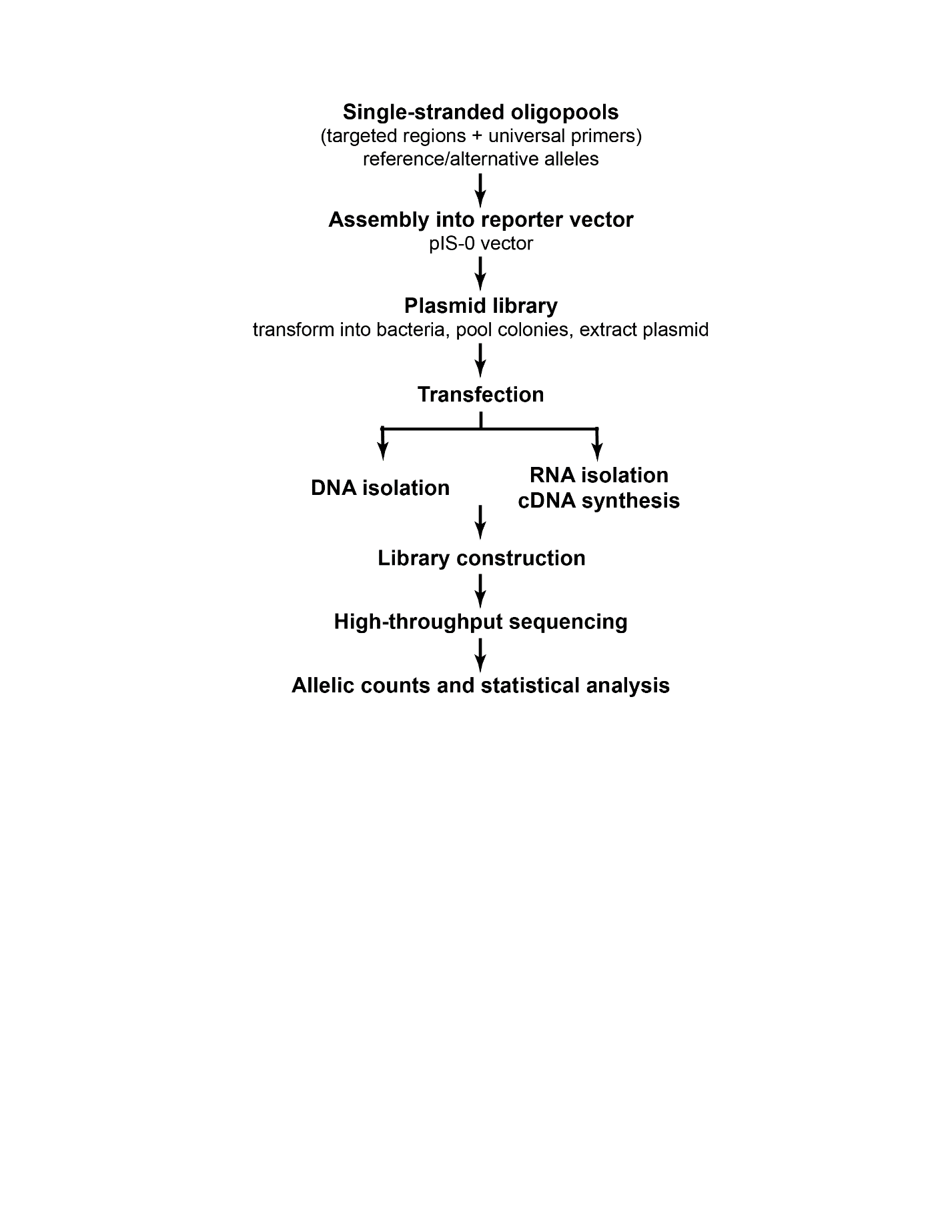

**Figure S3. Summary of experimental protocol.** A pool of oligonucleotides representing the reference and alternative alleles is printed and assembled into the pIS-0 reporter vector. The cloned oligo library is then transformed and expanded by bacterial colony growth to create the plasmid library. The plasmid library is transfected into the cell line of interest, and the resulting cellular DNA and RNA are isolated. The recovered cloned oligos are barcoded and sequenced. The counts of each oligo are used to calculate the effect of each variant (see Methods).

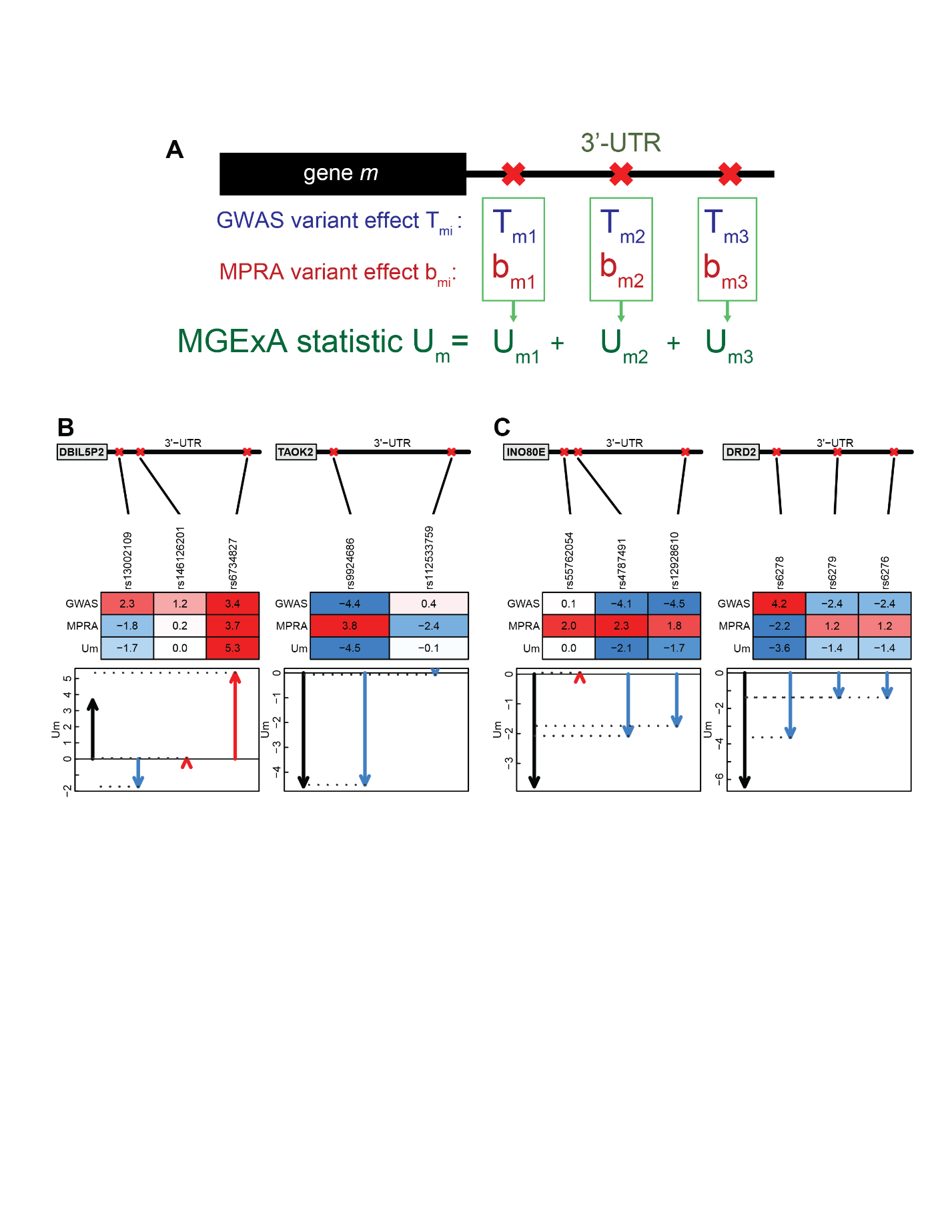

**Figure S4. MGExA identifies genes with 3’-UTRs associated with traits of interest by combining GWAS summary statistics with MPRA-derived variant effects.** (a) For each SNP $i$ in a gene’s 3’-UTR, the GWAS-derived effect sizes and MPRA-derived variant effects are multiplied and summed, yielding the $U_{mi}$ score (see Methods). For each gene, the MPRA effects of each MPRA-evaluated SNP are combined with the GWAS effects for a phenotype of interest, and the sum of these values equals $U_{m}$. (b-c) Schematic representations of MGExA evaluation for (b) DBILP2 and TAOK2 in SH-SY5Y and (c) INO80E and DRD2 in microglia cells.

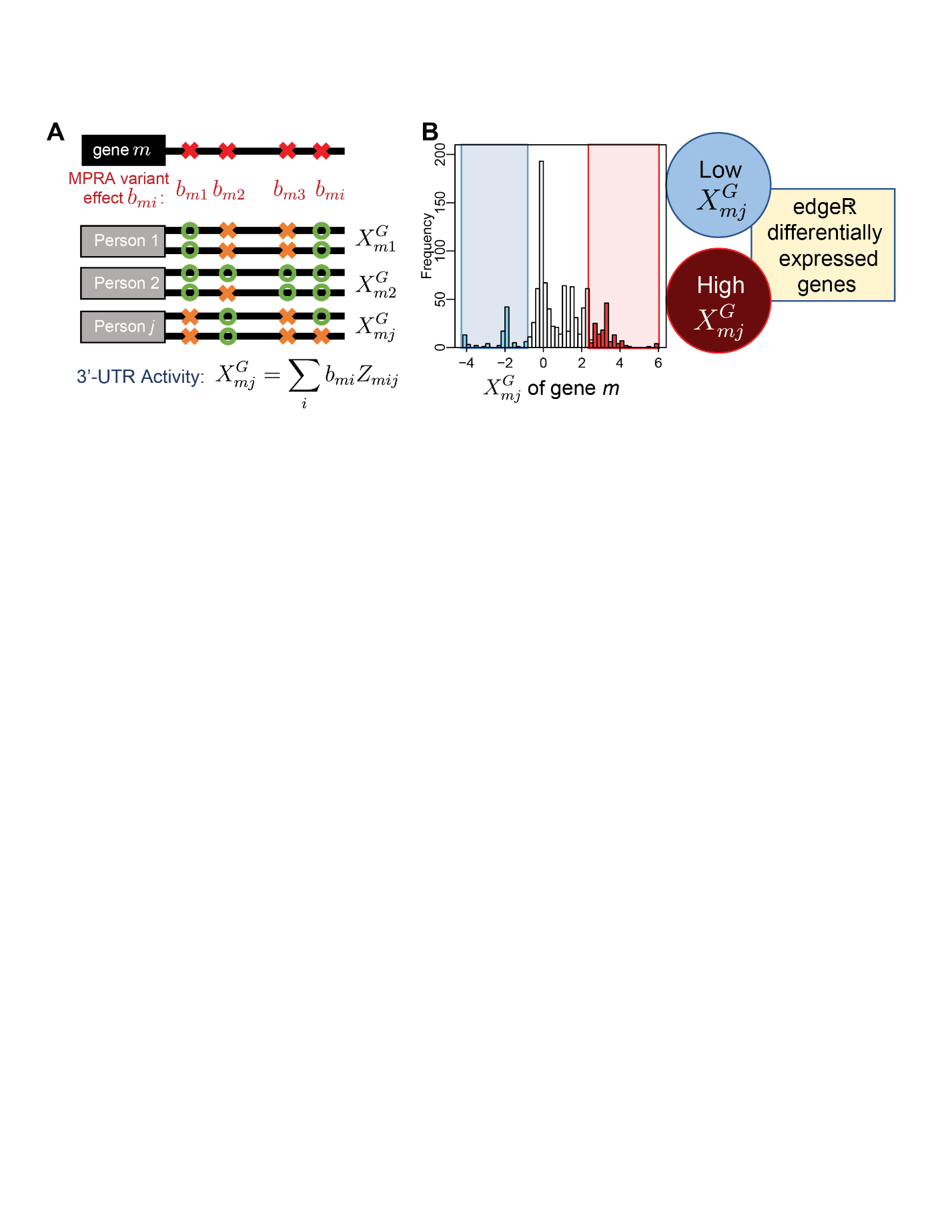

**Figure S5. Calculation of individual-level 3’-UTR activity for a gene to stratify individual samples into low and high 3’-UTR activity groups.** (a) For each individual, the genotype (dosage of the alternative allele) of each 3’-UTR SNP in a gene is multiplied by the MPRA-derived variant effect, and these values are summed across the 3’-UTR SNPs in the gene. (b) Then, this activity score is used to stratify samples into high and low activity groups consisting of the top and bottom 33% by effect score, and using RNA-seq data, the differentially expressed genes for this stratification of samples are identified by edgeR.

**Supplemental Tables**

**Table S1.** List of GWAS studies associated with neurological disorders from GWAS catalog used to identify lead SNPs.

**Table S2.** Lead SNPs from GWAS catalog used to identify candidate SNPs for MPRA. SNPs were those with p < 10^-5^ from GWAS of neurological disorders. SNPs for evaluation by MPRA were found in the regions around each lead SNP between the furthest SNPs on its left and right with r^2^ > 0.8.

**Table S3.** MPRA results from all 13,515 candidate SNPs in SH-SY5Y (SH in table) and microglia (MG) cells. Gene: gene symbol for 3’-UTR where SNP is located; rsID: ID of SNP from dbSNP v151; REF: reference allele; ALT: alternative allele; chr: chromosome; pos: SNP position in genome assembly GRCh38; lead SNPs: lead SNPs whose LD regions overlap the MPRA SNP; traits associated with lead SNPs: traits for GWAS that lead SNPs were associated with; SH: results derived from SH-SY5Y cells; MG: results derived from microglia cells; MPRA Variant Effect: the variant effect of the alternative allele in the MPRA; MPRA p-value: the p-value of the variant effect; MPRA FDR: false discovery rate; ALT frequency: the ALT allele counts divided by the sum of ALT and REF allele counts; mean count: the mean number of reads across 6 biological replicates for each allele and cell type.

**Table S4**. Heritability enrichment in alcoholic drinks per week and alcohol use disorder (AUD) GWAS for MPRA SNPs. Share of heritability in each GWAS was found for the set of SNPs that significantly alter (FDR < 0.05) gene expression in the MPRA of each cell line (SH-SY5Y and microglia) and that of the set of all MPRA SNPs. A permutation test was conducted to evaluate the significance of these differences by randomly generating 1000 sets of MPRA SNPs with the same number of SNPs as the set of FDR-significant SNPs and calculating the proportion with share of heritability greater than the share of heritability of the FDR-significant SNPs. SNP set share of heritability: the proportion of overall heritability of a trait explained by the selected SNPs, calculated by LDAK from GWAS summary statistics (see Methods); Expected Share of Heritability: the average heritability share for SNPs in the GWAS multiplied by the size of the SNP set; SNP Set Heritability Enrichment: the share of heritability divided by the expected share of heritability; Median share of heritability: the median heritability of the 1000 randomly generated SNP sets; Heritability Enrichment of MRPA FDR < 0.05 SNP Set Over Median: the share of heritability of the set of SNPs with MPRA FDR < 0.05 divided by the median share of heritability; fraction of random sets with heritability share larger than MPRA FDR < 0.05 set (p-value): the proportion of the 1000 randomly generated SNP sets with share of heritability larger than the set of SNPs with MPRA FDR < 0.05.

**Table S5.** Results of all genes evaluated by MGExA using GWAS summary statistics for GSCAN drinks per week phenotype. Um: the statistic derived from MGExA; p: the p-value associated with Um; FDR: the false discovery rate of p; SNPs: the number of SNPs used to evaluate each gene.

**Table S6**. A list of target sequences used for gene inhibition in Perturb-seq.

**Table S7**. Numbers of differentially expressed genes (DEGs) after knocking down each target gene using CRISPRi.

**Table S8**. Gene set enrichment results for effect of CRISPRi gene knock-down. Table includes target genes with successfully perturbed cells and KEGG pathways with adjusted p-value < 0.05.

**Table S9.** Significant genes and group sizes after stratifying individuals by 3’-UTR activity. Genes found by MGExA to be FDR < 0.2 in both cohorts of alcohol consumption GWAS or both cohorts of alcohol use disorder GWAS were used to stratify brain tissue samples by low and high 3’-UTR activity (see Methods). The 3’-UTR activity was calculated using SH-SY5Y MPRA results (SH genes) or microglia MPRA results (MG genes). Table shows the number of significant genes (FDR < 0.05) differentially expressed between the two groups as well as the number of individual samples in each group.

**Table S10.** Gene set enrichment results for genes differentially expressed in brain tissues stratified by imputed genetic component using MPRA results from SH-SY5Y cells. Table includes KEGG pathways with adjusted p-value < 0.05.

**Table S11.** Gene set enrichment results for genes differentially expressed in brain tissues stratified by imputed genetic component using MPRA results from microglia cells. Table includes KEGG pathways with adjusted p-value < 0.05.

**Table S12.** Primers used to barcode cDNA and plasmid DNA.

| **Name** | **Description** | **Sequence** |
| --- | --- | --- |
| HJ7211 | 1st strand - cDNA | ACA CGA CGC TCT TCC GAT CTN NGA AGA CTG NNN NNN NNN NCG GCC GCC CCG ACT CTA GAA CG |
| HJ7212 | 1st strand - plasmid DNA | ACA CGA CGC TCT TCC GAT CTN NNC ATT GCA CNN NNN NNN NNC GGC CGC CCC GAC TCT AGA ACG |
| HJ7214 | 2nd strand | CAG ACG TGT GCT CTT CCG ATC NAA TCC AGG CGC CGT GTA ATT CTA GGA GCT C |
| HJ7215 | 2nd strand | CAG ACG TGT GCT CTT CCG ATC NNT GAG GAG ACG CCG TGT AAT TCT AGG AGC TC |
| HJ7216 | 2nd strand | CAG ACG TGT GCT CTT CCG ATC NNN GAC TTG GAC GCC GTG TAA TTC TAG GAG CTC |
| HJ7217 | 2nd strand | CAG ACG TGT GCT CTT CCG ATC NTC TCA CCA CGC CGT GTA ATT CTA GGA GCT C |
| HJ7218 | 2nd strand | CAG ACG TGT GCT CTT CCG ATC NNG TGC GTT ACG CCG TGT AAT TCT AGG AGC TC |
| HJ7219 | 2nd strand | CAG ACG TGT GCT CTT CCG ATC NNN TCA TCG AGC GCC GTG TAA TTC TAG GAG CTC |
| HJ7220 | Passport-Seq-PCR | ACA CGA CGC TCT TCC GAT CT |
| HJ7221 | Passport-Seq-PCR | CAG ACG TGT GCT CTT CCG ATC |

**References**

1.	Buniello, A., MacArthur, J.A.L., Cerezo, M., Harris, L.W., Hayhurst, J., Malangone, C., McMahon, A., Morales, J., Mountjoy, E., Sollis, E., et al. (2019). The NHGRI-EBI GWAS Catalog of published genome-wide association studies, targeted arrays and summary statistics 2019. Nucleic Acids Res *47*, D1005-d1012. 10.1093/nar/gky1120.

2.	Lai, D., Wetherill, L., Bertelsen, S., Carey, C.E., Kamarajan, C., Kapoor, M., Meyers, J.L., Anokhin, A.P., Bennett, D.A., Bucholz, K.K., et al. (2019). Genome-wide association studies of alcohol dependence, DSM-IV criterion count and individual criteria. Genes Brain Behav *18*, e12579. 10.1111/gbb.12579.

3.	Sherry, S.T., Ward, M.H., Kholodov, M., Baker, J., Phan, L., Smigielski, E.M., and Sirotkin, K. (2001). dbSNP: the NCBI database of genetic variation. Nucleic Acids Res *29*, 308-311. 10.1093/nar/29.1.308.
